# Dissection of brain-wide spontaneous and functional somatosensory circuits by fMRI with optogenetic silencing

**DOI:** 10.1101/2021.07.22.453311

**Authors:** Won Beom Jung, Haiyan Jiang, Soohyun Lee, Seong-Gi Kim

**Author notes:** Corresponding author: Seong-Gi Kim, IBS Center for Neuroscience Imaging Research, N Center Sungkyunkwan University, Suwon 16419, Republic of Korea. **Author Contributions:** W.B.J. and S.G.K. conceived and planned the experiments; W.B.J. and H.J. performed the experiments; W.B.J. analyzed the data; S.L. provided critical discussion and insights; and W.B.J., S.L. and S.G.K. wrote the manuscript. **Competing Interest Statement:** The authors declare no competing interests.

## Abstract

To further advance functional magnetic resonance imaging (fMRI)-based brain science, it is critical to dissect fMRI activity at the circuit level. To achieve this goal, we combined brain-wide fMRI with neuronal silencing in well-defined regions. Since focal inactivation suppresses excitatory output to downstream pathways, intact input and suppressed output circuits can be separated. Highly specific cerebral blood volume-weighted fMRI was performed with optogenetic simulation of local GABAergic neurons in mouse somatosensory regions. Brain-wide spontaneous somatosensory networks were found mostly in ipsilateral cortical and subcortical areas, which differed from the bilateral homotopic connections commonly observed in resting-state fMRI data. The evoked fMRI responses to somatosensory stimulation in regions of the somatosensory network were successfully dissected, allowing the relative contributions of spinothalamic (ST), thalamocortical (TC), corticothalamic (CT), corticocortical (CC) inputs and local intracortical circuits to be determined. The ventral posterior thalamic nucleus (VPL) receives ST inputs, while the posterior medial thalamic nucleus (POm) receives CT inputs from the primary somatosensory cortex (S1) with TC inputs. The secondary somatosensory cortex (S2) receives mostly direct CC inputs from S1 and a few TC inputs from the VPL. The TC and CC input layers in cortical regions were identified by laminar-specific fMRI responses with a full width at half-maximum of <150 µm. Long-range synaptic inputs in cortical areas were amplified approximately 2-fold by local intracortical circuits, which is consistent with electrophysiological recordings. Overall, whole-brain fMRI with optogenetic inactivation revealed brain-wide, population-based long-range circuits, which could complement data typically collected in conventional microscopic functional circuit studies.

## Introduction

Functional magnetic resonance imaging (fMRI) has led to tremendous advances in brain science by enabling noninvasive mapping of both functional regions with various stimuli and resting-state functional connectivity. Evoked fMRI can be used to detect the strength of hemodynamic responses to external stimuli (1–3), whereas resting-state fMRI (rs-fMRI) measures the degree of synchrony between fMRI time series among anatomically distinct brain regions at rest (4–7). Both evoked and resting-state functional networks contain multiple brain regions that are hierarchically yet reciprocally connected by ascending thalamocortical (TC), descending corticothalamic (CT), corticocortical (CC) and intracortical (IC) circuits (8). Therefore, it is critical to determine the relative contributions of different circuits to fMRI findings to better understand brain functions and resting-state connectivity.

To identify the contributions of neural networks to fMRI responses, we aimed to silence neural activity in well-defined regions via temporally specific optogenetic control of a given neural population (9). Inhibiting one cortical region inevitably suppresses excitatory output to downstream signaling pathways (e.g., CT, CC, and IC pathways) (10–12), and the downregulated neuronal activity reflects the degree of interregional communication under basal conditions (12). Therefore, fMRI with cortical inactivation is beneficial for brain-wide mapping of functional connectivity in the resting state. Similarly, local silencing during exposure to external stimuli suppresses downstream activity (output-related circuits) without compromising the upstream and/or collateral inputs from other brain regions (input-related circuits) (11, 13–16); thus, downstream circuit contributions to fMRI can be determined by comparing evoked fMRI responses with and without focal inactivation.

In the current study, we adopted a widely investigated somatosensory network involving multiple brain regions, such as the first-order ventral posterior thalamic nucleus, higher-order posterior medial thalamic nucleus (POm), primary somatosensory cortex (S1), secondary somatosensory cortex (S2), and primary motor cortex (M1) (17, 18). Anatomical tracing studies have demonstrated the existence of complex monosynaptic networks across broad somatosensory-related brain areas (19–21). In addition, microscopic functional circuits in preselected areas have been delineated using electrophysiology (22–26) and optical recordings (27, 28) combined with optogenetic and chemogenetic tools. Recently, macroscopic functional activity in the whole brain was mapped by fMRI (29–32). Thus, the somatosensory system is an ideal model for investigating whether the contributions of different projections to fMRI responses can be separated.

To investigate spontaneous and sensory-evoked fMRI signals at the circuit level, we combined highly specific cerebral blood volume (CBV)-weighted fMRI at an ultrahigh magnetic field strength of 15.2 T with optogenetic stimulation of local GABAergic neurons using the vesicular GABA transporter (VGAT)-channelrhodopsin-2 (ChR2)-enhanced yellow fluorescent protein (EYFP) transgenic mouse line (33). We examined how different cortical areas, including the primary somatosensory forelimb (S1FL), M1 and S2, affect activity in cortical and subcortical brain regions to determine causal relationships in the information flow among multiple brain areas.

## Materials and Methods

### Animal preparation

A total of 45 mice (21.4–31.5 g, 6–13 weeks old, male/female = 36/9) were used in six different studies: 1) 22 transgenic mice for fMRI with cortical inactivation (VGAT-ChR2-EYFP, n=8 for S1FL, n=7 for M1, and n=7 for S2), 2) 6 VGAT-ChR2 mice for electrophysiology recording, 3) 3 naïve mice for contrast agent dose-dependent MRI to optimize the CBV-weighted fMRI protocol (C57BL/6; Orient Bio, Seongnam, Korea), 4) 3 naïve mice for the fMRI study on light-induced heating effects, 5) 5 naïve and 4 VGAT-ChR2 mice for conventional resting-state CBV-weighted fMRI, and 6) 1 VGAT-ChR2 positive mouse and 1 VGAT-ChR2 negative littermate for histological confirmation of GABAergic interneuron-specific expressions of ChR2 in transgenic VGAT-ChR2-EYFP mice. The VGAT-ChR2-EYFP mice (B6.Cg-Tg (Slc32a1-COP4*H134R/EYFP) 8Gfng/J) were bred in-house from breeding pairs obtained from Jackson Laboratory (see Fig. S1; Bar Harbor, ME, USA). Animals were housed in independently ventilated cages under controlled temperature and humidity conditions and a 12-hour dark-light cycle.

A stereotactic surgical procedure (34) was performed to implant fiber-optic cannulas (Thorlabs, Newton, New Jersey, USA) in mice (VGAT-ChR2-EYFP, n=22; C57BL/6, n=3) at 6-7 weeks of age for optogenetic fMRI. In short, the mice were anesthetized by an intraperitoneal (IP) bolus injection of a ketamine and xylazine mixture (100 mg/kg and 10 mg/kg, respectively) and fixed in a stereotaxic frame (SR-AM, Narishige, Tokyo, Japan) for surgery. Meloxicam (1 mg/kg) was administered subcutaneously to provide pain relief and reduce inflammation. After an incision was made on the animal’s scalp, the skull was completely washed and dried. The skull was then thinned over the target area to make a burr hole (∼ 100 µm diameter). The optical fiber cannula (105 µm inner core diameter, NA = 0.22) was then inserted slowly into the right S1FL (AP: −0.2 mm relative to bregma, ML: + 2.2 mm and DV: + 0.5 mm relative to the surface), M1 (AP: + 0.05 mm, ML: + 1.1 mm and DV: + 0.25 mm) and S2 (AP: - 1.1 mm, ML: + 4.2 mm and DV: + 0.9 mm) areas (see Fig. S2A for visualization of tip position) through the burr hole at a rate of 1 µm/sec via a micromanipulator (SMX-model, Sensapex, Oulu, Finland). With the fiber-optic implant in place, a biocompatible silicone elastomer (Kwik-Sil, World Precision Instruments, Sarasota, FL, USA) was applied to enclose the fiber implantation site, and dental cement (SB, Sun-Medical Co., Shiga, Japan) was then applied to thinly cover the area around the fiber cannula to fix it onto the skull. After a recovery period of at least 2 weeks, mice underwent optogenetic fMRI experiments.

The functional experimental procedure was previously described in detail (30, 35). In short, mice were anesthetized using a mixture of ketamine (Yuhan, Korea) and xylazine (Rompun^Ⓡ^, Bayer, Korea) (100/10 mg/kg for initial IP, and 25/1.25 mg/kg intermittent IP injections every 45-50 min) under self-breathing through a nose cone that provided a continuous supply of oxygen and air (1:4 ratio) at a rate of 1 liter/min (SAR-1000, CWE, Ardmore, USA or TOPO, Kent Scientific Corporation, Torrington, CT, USA) to maintain an oxygen saturation level > 90% (36). During the experiments, electrocardiograms and motion-sensitive respiratory signals were continuously monitored (Model 1030, Small Animal Instruments Inc., Stony Brook, USA for fMRI experiments and PhysioSuite, Kent Scientific Corp, USA for electrophysiology studies). Also, the body temperature of the animals was maintained at 37 ± 0.5 °C with a warm-water heating system and a rectal thermometer.

### Stimulation

The stimulation parameters were controlled by a pulse generator (Master 9; World Precision Instruments, Sarasota, FL, USA). For light stimulation, blue light was delivered to the target cortex via a fiber-optic cable coupled to a 473-nm diode-pumped solid-state laser (MBL-III-473, Changchun New Industries Optoelectronics Tech. Co., Ltd, Changchun, China). The constant output power was calibrated to be 3 mW at the tip of the optic fiber, measured by a power meter (PM100D, Thorlabs, USA). Light stimulation was then applied at 20 Hz with a pulse width of 10 ms via a 105 μm diameter fiber. This experimental setting leads to a time-averaged light power of 69.3 mW/mm^2^, which is below the range at which light-induced artifactual responses are generated (37, 38). To block undesired activation of the visual pathway by light leakage inside the magnet bore, the connection between the fiber-optic cable and the implanted cannula was covered with heat-shrinkable sleeves, and the eyes of the animals were covered with a biocompatible silicone elastomer. For somatosensory stimulation, the left forepaws of the mice were electrically stimulated. A pair of needle electrodes (30 G) were inserted under the plantar skin of the forepaws, and electrical pulse stimuli with a current intensity of 0.5 mA (ISO-Flex, AMPI, Jerusalem, Israel), pulse width of 0.5 ms, and frequency of 4 Hz were triggered by a pulse generator (35). Each functional trial consisted of a 60-s (30 s for electrophysiology) prestimulus, 20-s stimulus, 60-s interstimulus interval, 20-s stimulus, and 60-s (30 s for electrophysiology) poststimulus period.

### Electrophysiology

Multiunit activity (MUA) and local field potential (LFP) were measured in the S1FL to confirm the suppression of neuronal activity during optogenetic cortical inactivation and to investigate the neural source of fMRI findings concerning long-range TC input and local recurrent activity (VGAT-ChR2-EYFP, n=6). The head was fixed in a stereotaxic frame (SR-10R-HT, Narishige, Tokyo, Japan), and the scalp was removed to expose the skull. After cleaning the skull, two holes (> 0.3 mm in diameter) were made with a dental drill to implant the stainless-steel screws for the ground and reference wires. Craniotomy was performed over the right somatosensory area, with a diameter of ∼2 mm. CBV-weighted optical imaging (Imager 3001, Optical Imaging Ltd., Rehovot, Israel) of the responses to left forepaw stimulation (FP) was performed to precisely determine the S1FL area. To simultaneously record neural activity during optogenetic stimulation, a 16-channel optoelectrode with 50 µm spacing between channels (A1×16-5 mm-50-177-OA16LP, NeuroNexus, Ann Arbor, MI, USA) was placed at the center of the predefined S1FL and slowly inserted at a speed of 1 μm/sec. The fiber tip in this optoelectrode was positioned ∼ 200 μm above the proximal recording site nearest to the cortical surface. In one mouse, a separate optic fiber was inserted into the middle of the cortex obliquely near the optoelectrode to determine optogenetic cortical inactivation across all cortical depths (see Fig. S2B-i for visualization of the tip position). During insertion, the electrode was held for 5 min every 100 µm and for 10 min when the tip of the electrode reached a cortical depth of ∼1 mm to stabilize the position. Electrophysiological signals were measured at a sampling rate of 30 kHz using a Blackrock Cerebus recording system (Blackrock Microsystems, Salt Lake City, Utah). Functional studies were interleaved across 3 different stimulus paradigms, and 5-8 trials were performed for each experimental condition for each animal.

### MRI experiments

All MRI experiments were performed on a 15.2 T magnetic resonance imaging (MRI) scanner with an actively shielded 6-cm diameter gradient operating with a maximum strength of 100 G/cm and a rise time of 110 μs (Bruker BioSpec, Billerica, MA, USA). A 15 mm ID customized surface coil for radiofrequency (RF) transmission/reception was centered on the imaging slices covering the somatosensory cortex, and the brain was positioned close to the isocenter of the magnet. Magnetic field homogeneity was optimized by global shimming, followed by the FASTMAP shimming protocol on the ellipsoidal volume covering the cerebrum (ParaVision 6, Bruker BioSpin).

#### Calibration of the amount of contrast agent used for CBV-weighted fMRI studies

Gradient-echo (GE) echo planar imaging (EPI), commonly used in functional studies, is sensitive to susceptibility artifacts arising from implanted optic fibers, particularly at long TEs and high field strengths (Fig. S3A). Importantly, the BOLD contrast measured by GE-EPI is highly sensitive to draining veins, resulting in poor specificity for cortical layers. CBV-weighted fMRI with a superparamagnetic contrast agent is an alternative method to GE-BOLD to improve the functional sensitivity and spatial specificity at neuronal active sites (39). CBV-weighted fMRI with a short TE has the advantage of reducing nonspecific BOLD contributions and minimizing susceptibility artifacts. Initially, we measured the relationship between contrast doses and *T*_2_^∗^ values by repeated injections of monocrystalline iron oxide nanoparticles (MIONs; Feraheme, AMAG Pharmaceuticals, Waltham, USA) into the tail vein until the cumulative dose reached 45 mg/kg (from 2 to 45 mg/kg) in 3 naïve mice (Fig. S3B). *T*_2_^∗^-weighted images were obtained using the multi-gradient-recalled echo (Multi-GRE) sequence with sixteen echo times = 1.5 to 18 ms and an interval of 1.1 ms, repetition time (TR) = 0.8 s, field of view (FOV) = 15 (read-out, x-axis) × 7.5 (phase-encoding, y-axis) mm^2^, matrix = 96 × 48, spatial resolution = 156 × 156 × 500 μm^3^, flip angle = 50°, 6 contiguous coronal slices and number of excitations (NEX) = 4.

#### fMRI combined with optogenetics

Our imaging strategy was to use a very short TE for fMRI data without susceptibility artifacts while maximizing the functional sensitivity. Therefore, TE was set to 3 ms for matching to tissue *T*_2_^∗^ (3.44 ± 0.08 ms in Fig. S3B) values, with MION at a 45 mg/kg dose. CBV-weighted fMRI data were acquired using the fast low-angle shot magnetic resonance imaging (FLASH) sequence with the following parameters: TR/TE = 50/3 ms, flip angle = 15°, FOV = 15 × 7.5 mm^2^, matrix size = 96 × 48, partial Fourier = 1.2 along the phase-encoding direction, spatial resolution = 156 × 156 × 500 μm^3^, 6 contiguous coronal slices and temporal resolution = 2 s. From individual transgenic mice (n=22 mice), fMRI data were acquired under three stimulus conditions: optogenetic cortical silencing (n=8 for S1FL targeting, n=7 for M1 targeting, and n=7 for S2 targeting), FP, and simultaneous silencing and FP. Data for the three stimulus conditions were acquired in an interleaved manner, with an interscan interval longer than 3 min. To address the potential heating-related artifacts induced by light, functional studies in naïve mice (n=3) were interleaved between optogenetic and FPs. For each stimulus condition, 15 fMRI trials were performed for each animal.

#### rs-fMRI

Conventional rs-fMRI, which is presumably related to the intrinsic network of spontaneous activity, was obtained to examine the resting-state somatosensory network. Ten 10-min rs-fMRI scans (i.e., 300 volumes) were obtained from two separate animal groups, naïve animals (n=5, twice each) and transgenic VGAT-ChR2 animals (n=4, twice each) with the same imaging parameters used for fMRI, after a few somatosensory fMRI scans to ensure the physiological condition of the mice responding to external stimuli under anesthesia.

### Histology

To confirm GABAergic interneuron-specific expression of ChR2 in home-bred transgenic mice, we performed histology of one VGAT-ChR2-EYFP positive mouse and one VGAT-ChR2-EYFP negative littermate mouse. Animals were anesthetized with ketamine/xylazine and intracardially perfused with saline, followed by 4% paraformaldehyde (PFA) for fixation. The brains were postfixed in 4% PFA overnight at 4 °C and then stabilized with 30% sucrose with 0.1% sodium azide solution at 4 °C for 3 days. The brains were coronally sectioned at a 40-μm thickness with a cryostat (CM 1950, Leica Biosystems). Two sections were selected to check ChR2 expression: one included the S1FL to validate the expression of ChR2 in cortical GABAergic neurons, and the other included the GABAergic neuron-rich thalamic reticular nucleus (TRN). The sections were incubated with mouse anti-GAD67 primary antibodies (1:300; Millipore) and decorated with secondary antibodies conjugated with Alexa 568 (1:350; Invitrogen) to label interneurons. Colocalization of GAD67-positive neurons with EYFP expression in transgenic mouse indicated that ChR2 was specific to GABAergic neurons. Nuclear counterstaining was also performed with DAPI solution (1:10,000; Sigma) in PBS. The stained brain slices were imaged with confocal laser microscopy (TCS SP8, Leica Microsystems). The green-, red-, and blue-colored representations depict the relative spectral weights for ChR2-EYFP-, GAD67-and DAPI-positive expression, respectively.

### Data analysis

#### Electrophysiology data analysis

To detect spiking activity, the extracellular recording traces were preprocessed as follows: first, raw data were high-pass filtered (> 300 Hz; 4^th^-order Butterworth filter); second, the common noise level was temporally removed by subtracting the trimmed mean (50%) filtered signals over all channels at each time point from each channel; third, MUA was quantified over 1 ms temporal bins by counting the number of spikes exceeding a threshold of 5 × the median absolute deviation in 30 s of prestimulus baseline data (40). The LFP data were extracted from the extracellular recording traces by bandpass filtering at 1 to 280 Hz and downsampled to 1 kHz. For the comparison between trials with FP at 4 Hz and optogenetic silencing at 20 Hz, spike trains and LFP waveforms during stimulation were aligned with a 250-ms moving window after the first stimulus onset of each block, and repeated runs (20 s × 2 blocks × 4 windows/s × # of trials) were then averaged (Fig. S8B).

To confirm the suppression of spontaneous neural activity during optogenetic cortical stimulation, each light pulse-induced putative inhibitory neural activity was excluded for the 10 ms light pulse period and followed up for 10 ms, resulting in spontaneous spike activity during the 30 ms period following each 20 Hz stimulation pulse (41). The firing rates were also calculated across the upper 8 and lower 8 channels to measure depth dependency.

To investigate the cortical depth-dependent MUA and LFP responses, we first performed inversion current source density (iCSD) analysis with the LFP waveform using the CSDplotter toolbox (42) to identify the location of L4. The synaptic inputs induced by FP were observed as a strong sink at depths of 0.4-0.56 mm, whereas the current sink generated by each optogenetic stimulation was shown at depths of 0.2-0.4 mm (Fig. S8C). Based on these laminar CSD profiles, the cortical depth of the S1FL was assigned as L2/3, L4 and L5/6 (Fig. S8D and S8E).

We compared the spiking responses and field potential observed during optogenetic silencing without and with FP. Since somatosensory-evoked recurrent excitatory circuits were suppressed by optogenetically activating inhibitory neurons in the S1FL, and the remaining activity indicated direct feedforward inputs from the thalamus (Diff_S1FL_). The contribution of silenced local recurrent circuits was calculated by the difference between TC input-driven activity and the total somatosensory-evoked response. To alleviate minor response timing errors from the calculation, three sensory-evoked MUA traces and LFP waveforms (TC input-driven activity vs. local recurrent activity vs. total activity) were fitted using a single Gaussian function and double gamma variate function, respectively, averaged across all channels and for each animal.

In the fitted MUA curve, activity parameters, including peak amplitude and the area under curve (AUC), and dynamic parameters, including onset time, peak time, and full-width at half-maximum (FWHM), were quantified. Response onset was defined as the first bin of five continuous bins in which the spike rate differed significantly from that observed at baseline activity (p < 0.05, one-sample t-test). In LFPs, the same parameters described for MUA were calculated except for the onset time.

#### Calibration of the amount of contrast agent

To determine MION dose-dependent tissue *T*_2_^∗^, multi-GRE data before and after every injection were fitted by a monoexponential function as 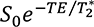, where *S*_0_ is the signal at TE = 0 ms. For quantification of tissue *T*_2_^∗^ changes, the ROI was defined as a two-dimensional (2D) sphere with a radius of 500 μm centered on the anatomical location of the S1FL with reference to the Allen mouse brain atlas.

#### Overall fMRI data analysis

All MRI data were analyzed with the Analysis of Functional Neuroimages package (AFNI) (43), FMRIB Software Library (FSL) (44), Advanced Normalization Tools (ANTs) (45), and MATLAB codes (MathWorks, Natick, USA).

To calculate group-averaged fMRI maps, a study-specific mouse brain template was constructed from CBV-weighted fMRI images based on a previously used pipeline (46, 47) as follows. First, the brain area was semiautomatically extracted from the temporal average of all fMRI images for each animal, and intensity nonuniformity was corrected with bias field estimation. Second, the individual fMRI images were aligned with each other so that the most representative image requiring the least transformation within the population could be selected (44). Third, all the images were spatially normalized into that representative image using both linear and nonlinear transformation and averaged to generate the mouse brain template. To identify the anatomical location, the study-specific template was coregistered with the Allen mouse brain atlas (19).

#### Generation of evoked-fMRI maps

The fMRI response maps for individual animals were generated using preprocessing and general linear model (GLM) analysis. The preprocessing steps were previously described in detail (30, 46). In short, preprocessing included slice timing correction, image realignment for minor head motions, linear signal detrending for signal drift removal, and time course normalization by the average of the baseline period. All repeated fMRI trials per stimulus condition for each animal were averaged and spatially coregistered to study-specific brain templates. Then, spatial smoothing was applied with a Gaussian kernel of one-pixel FWHM to minimize potential misalignment and to enhance functional detectability. Individual fMRI maps were calculated by a GLM analysis with design matrices where two 20-s stimulus blocks were convolved by the data-driven hemodynamic response function (HRF). The HRF was determined from the animal-wise averaged time course in the S1FL during FP by fitting with a two-gamma variate function. After GLM analysis, group-averaged fMRI maps of FP, optogenetic stimulation only (Opto), and combined forepaw and optogenetic stimulation (Comb) were generated for the S1FL, M1 and S2 silencing conditions with a one-sample t-test using nonparametric inference (44), with significance at threshold-free cluster enhancement (TFCE)-corrected p < 0.05. Standardized regression coefficients (β value) were mapped.

To determine somatosensory input-driven responses without the feedforward or local contribution from the inactivation site, fMRI time courses during optogenetic cortical inactivation with and without FP were subtracted (Comb – Opto), and difference maps (Diff) were obtained with a significance level of uncorrected p < 0.01 (paired t-test). In addition, all animal data (n=22) were used to generate group-averaged FP activation maps. The group-averaged fMRI maps were overlaid on the study-specific brain template, and the fMRI active sites were determined according to the Allen mouse brain atlas.

#### Quantitative ROI analyses for circuit analysis

For quantitative analysis, five different somatosensory ROIs corresponding to active sites during FP were defined based on the Allen mouse brain atlas: S1FL, M1, S2, ventral posterolateral nucleus (VPL), and POm. Time courses from each ROI were obtained for FP, Opto, Comb, and Diff conditions in each animal, and fMRI signal changes were calculated by averaging responses over the stimulus condition, excluding the initial 6 s after stimulus onset to obtain steady-state positive or negative responses without the initial transition period. An increase in CBV increased the amount of iron oxides within the voxel, consequently decreasing fMRI signals. Thus, the original CBV-weighted fMRI signal changes were inverted for better visualization in all analyses.

#### Seed-based rs-fMRI mapping

For rs-fMRI data without any stimulations, preprocessing steps similar to those used for evoked fMRI were included: slice timing correction, image realignment, linear detrending, and voxel-wise time course normalization. To stabilize functional connectivity, additional preprocessing steps were applied: spiking signal removal, data cleaning by regressing out nuisance variables including 12 motion confounds (6 parameters + temporal derivatives) and brain global signals, bandpass filtering (0.01 < f < 0.2 Hz) (35) and spatial smoothing with a Gaussian kernel of a 0.5 mm FWHM. Three seed ROIs of S1FL, M1 and S2 in the right hemisphere were anatomically defined based on the Allen mouse brain atlas. To identify the resting-state connectivity dependent on mouse strain, rs-fMRI data from 5 naïve mice (Fig. 2B) and from 4 transgenic mice (Fig. S4A) were separately analyzed. In common space, functional connectivity maps for individual animals were calculated by correlation analysis of the mean time courses of each seed ROI and voxel-wise time series in the whole brain. To generate group-averaged rs-fMRI maps, voxel-wise correlation coefficient (r) was converted to a normally distributed z-score using Fisher’s r-to-z transformation, and a one-sample t-test was performed (TFCE-corrected p < 0.05). After statistical testing, the results were inversely transformed back to correlation coefficients to represent group-averaged rs-fMRI maps.

#### Resting-state connectivity analysis

To investigate spatial patterns of spontaneous somatosensory network, we generated the two different connectivity maps with 1) cross-correlation coefficients (r) of rs-fMRI using temporal correlation in S1FL, M1, and S2 seed-regions (see subsection of *seed-based rs-fMRI mapping*) and 2) regression coefficients (β value) of fMRI with cortical silencing in S1FL, M1, and S2 areas using GLM analysis (Opto) (see subsection of *generation of evoked-fMRI maps*). The degree of rs-fMRI correlation between two different regions often considers as a proxy to spontaneous neural network strength (6), while fMRI signal changes in the networked areas causally induced by cortical silencing is related to the degree of interregional communication with source area under basal conditions (12). To compare the relative strength of spontaneous functional connectivities measured by rs-fMRI and cortical silencing fMRI (Opto), 1) the magnitude of functional connectivity (z-scores by Fisher’s r-to-z transform) within predefined somatosensory ROIs was normalized to the strongest connectivity with a seed area (given a maximum z-value equal to 1) in rs-fMRI data of naïve mice, 2) whereas the fMRI change in spontaneous connectivity during cortical silencing fMRI (Opto) was normalized to the response at the optogenetic stimulation site in VGAT-ChR2 mice. To examine which functional connectivity patterns closely reflect the intrinsic neural networks, tracer-based connectivity maps for S1FL (experiment #112229814), M1 (experiment #100141563) and S2 injection (experiment #112514915) were obtained from the Allen Institute (19). The connectivity strengths in networked areas were normalized by the projection density in the injection site (Fig. S4B).

#### Layer analysis of fMRI responses

To measure synaptic input layer-specific fMRI responses, all aforementioned preprocessing steps except spatial smoothing were applied. Cortical areas, including the S1FL, M1, and S2, were flattened by radially projecting 22 lines perpendicular to the cortical edges (48). The cortical depth profiles were resampled to double using bicubic interpolation, leading to a nominal resolution of 78 µm. Then, in each animal, an averaged profile was obtained in the S1FL and M1 by averaging 5-pixel lines (0.78 mm width) and in S2 by averaging 10-pixel lines (1.56 mm width). Group-averaged percent change maps were calculated from the percent change maps of individual animals. The fMRI signal changes within the same cortical depth were calculated in the same manner described above. Laminar boundaries for each cortex were defined as a cortical thickness distribution based on the Allen mouse brain atlas.

#### Statistics

All quantitative values are presented as the mean ± standard error of the mean (SEM). For a comparison of the suppression of spontaneous MUA firing rates with and without optogenetic inactivation (Fig. 1B-ii), an evaluation of fMRI signal changes for measuring heating-related fMRI artifacts (Fig. 1D), a comparison of fMRI signal changes between long-range sensory inputs (i.e., difference between cortical inactivation with and without FP) and total sensory-evoked activity (FP only) (Fig. 3C, 3D, 4C, 4D and 6B), and the analysis of the relative fraction of long-range inputs and local recurrent activities in the MUA and LFP (Fig. 4G), a paired t-test was conducted to determine the statistical significance of differences. For a comparison of fMRI signal changes across somatosensory fMRI experiments in different animal groups, one-way analysis of variance (ANOVA) was conducted (Fig. 3C, 3D, 4C, 4D and 6B). To compare the cortical depth-dependent MUA and LFP data (Fig. S8F-H), one-way ANOVA with repeated measures was performed. All ANOVA tests were followed by the Bonferroni *post hoc* test for multiple comparisons. Statistical significance was considered at p < 0.05.

**Figure 1.**
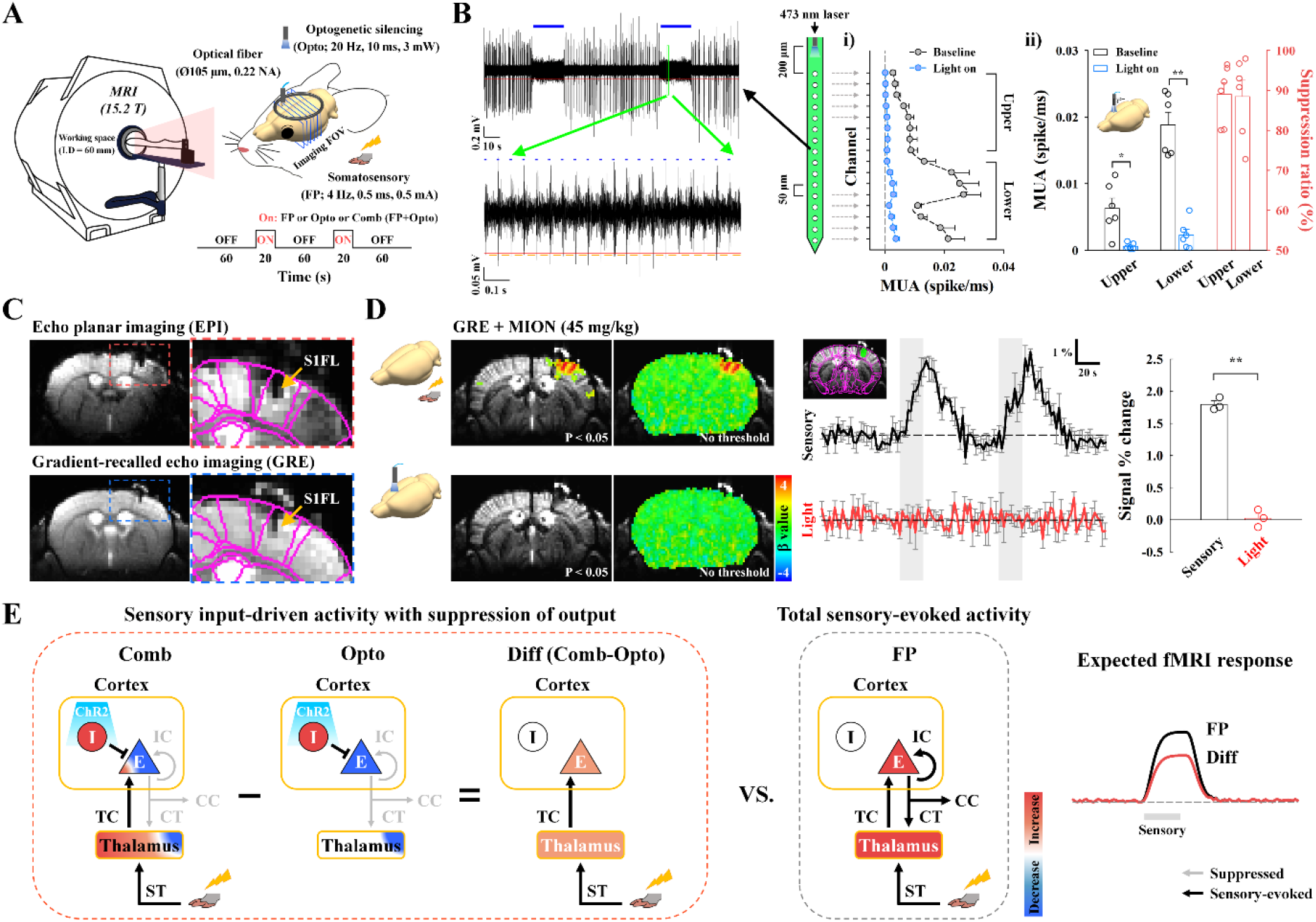
Experimental setup for high-resolution CBV-weighted optogenetic mouse fMRI and conceptual neural circuit analysis. **(A)** Schematic of multislice fMRI at 15.2 T during forepaw somatosensory stimulation (FP) and during cortical inactivation without (Opto) and with (Comb) somatosensory stimulation. **(B)** Silencing spontaneous activity of excitatory neurons in the S1FL by optogenetic excitation of inhibitory neurons in VGAT-ChR2 mice (n=6). A 16-channel optoelectrode with 50 µm interchannel spacing was inserted to a 1 mm depth, and photostimulation was delivered on the surface of the cortex. During two 20-s photostimulation periods (blue bars in the left example recording trace), MUAs were reduced; in the expanded view, MUA is shown to slightly increased during each pulse (blue dots) due to enhanced inhibitory activity but was near zero between pulses (yellow lines) due to suppressed spontaneous activity. During 20 s of photostimulation, spontaneous activity in i) all cortical layers (200–1000 µm depth) and ii) both the upper and lower 8 channels was mostly suppressed by activation of inhibitory neurons. Similar uniform suppression across all cortical depths was achieved during photostimulation at the middle of the cortex (see Fig. S2B). Blue horizontal bar, 20-s optogenetic stimulus; red, MUA amplitude threshold; yellow, spontaneous MUA count duration; error bar, SEM; *p < 0.05 and **p < 0.01 (paired t-test); suppression ratio in panel ii, attenuation rate of neural activity during optogenetic silencing relative to the baseline activity. **(C)** High-resolution MRI images of the brain with an optical fiber targeting the S1FL. The image artifacts of distortion and signal drops caused by the optical fiber in gradient-echo EPI (red-dashed box) were minimized by the adoption of gradient-echo imaging with a short echo time of 3 ms (yellow arrow, fiber position). For the CBV-weighted fMRI study, a 45 mg/kg dose of a superparamagnetic monocrystalline iron oxide nanoparticle (MION) agent was injected into the animals’ blood. S1FL, primary somatosensory area of the forelimb. Images of a fiber position and a choice of the MION dose are shown in Fig. S3. **(D)** No artifactual fMRI responses to light stimulation were observed in naïve mice. To assess potential light-induced MRI artifacts, functional studies in naïve mice were conducted with interleaved forepaw stimulation and photostimulation of the S1FL with the same experimental protocol used for fMRI with cortical inactivation (n=3). Somatosensory fMRI was used as the internal control to ensure the reliability of fMRI responses to external stimuli. Sensory-evoked fMRI responses were observed in the S1FL, but responses during photostimulation were absent in fMRI maps with and without a statistical threshold (uncorrected p < 0.05) and in time courses of the S1FL. Thus, fMRI signal changes resulting from light-induced tissue heating were negligible. Black and red time traces, somatosensory-evoked and photostimulated fMRI time courses in the S1FL, respectively; gray vertical bar in time courses, 20-s stimulus; error bars, SEM; **p < 0.01 (paired t-test). **(E)** Schematic diagrams to dissect sensory-evoked long-range and local activity by fMRI with the assistance of optogenetic cortical inhibition. In a simplified circuit, excitatory neurons (E) interacting with inhibitory interneurons (I) in the cortex received thalamocortical inputs from the thalamus and produced spiking outputs for intracortical (IC), corticocortical (CC) and corticothalamic (CT) downstream activity. Silencing cortical excitatory neurons by activating ChR2-expressing cortical inhibitory neurons suppressed outputs to downstream CT, CC and IC pathways (Opto), while simultaneous forepaw stimulation induced sensory-evoked upstream ST and TC inputs to the thalamus and cortex, respectively (Comb). After subtracting fMRI responses to optogenetic inactivation without forepaw stimulation (Opto) from those with forepaw stimulation (Comb), the difference signal was related to upstream input-driven responses (Diff; Comb-Opto) without the direct contribution of inhibitory neural activity. The relative contribution of downstream CT, CC and IC pathways can be determined from the total sensory-evoked activity (Diff vs. FP). E, excitatory cell; I, inhibitory cell; ST, spinothalamic pathway; TC, thalamocortical pathway; CT, corticothalamic pathway; CC, corticocortical pathway; IC, intracortical pathway; blue-to-red color scale, decrease-to-increase relative to baseline activity.

**Figure 2.**
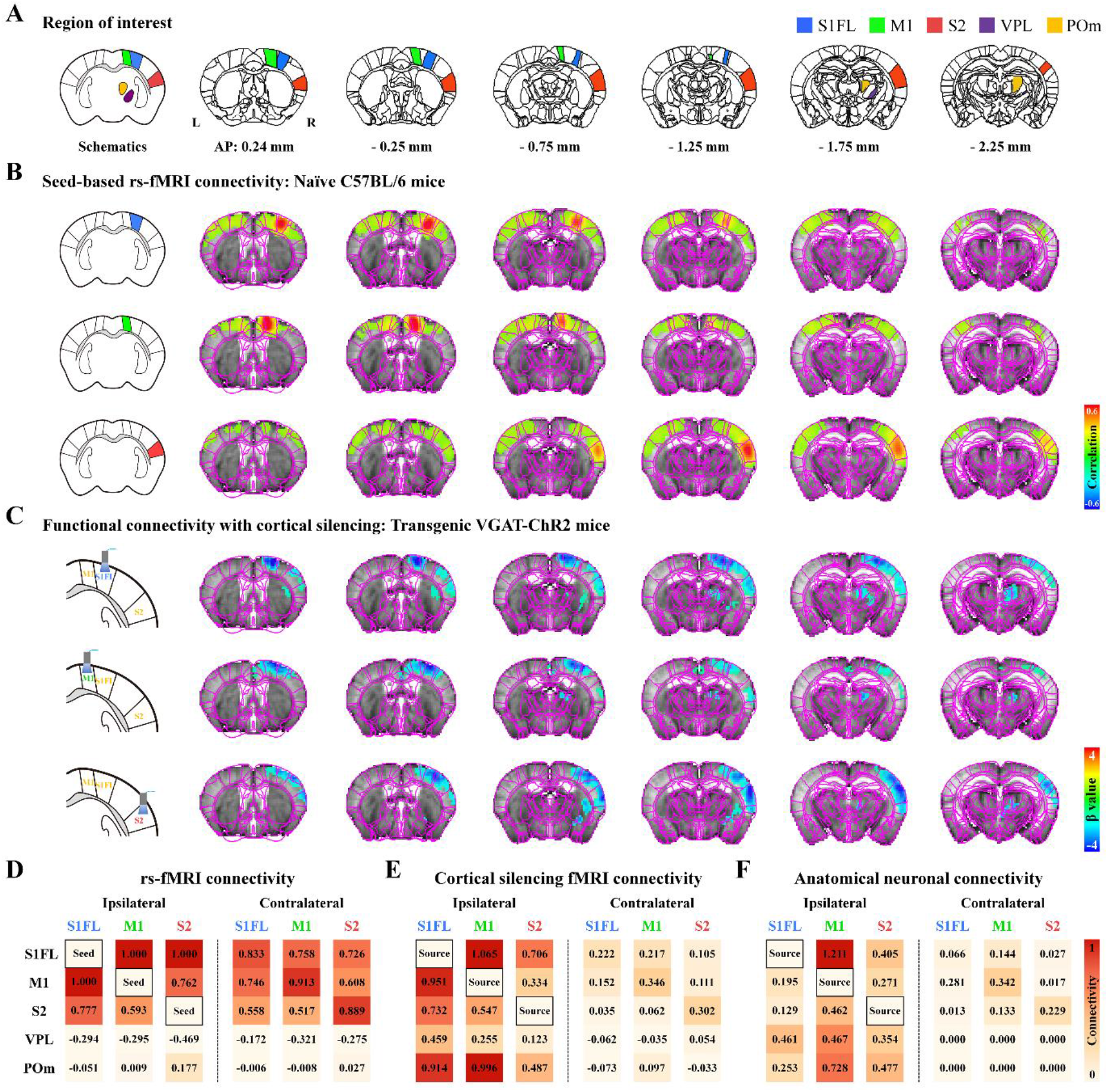
The spontaneous resting-state somatosensory network with optogenetic neuronal inactivation differs from conventional resting-state fMRI connectivity. **(A)** Allen mouse brain atlas-based anatomic ROIs in the somatosensory network were chosen for analyzing resting-state connectivity. S1FL, primary somatosensory area of the forelimb; M1, primary motor area; S2, secondary somatosensory area; VPL, ventral posterolateral nucleus; POm, posterior medial nucleus. **(B-C)** Brain-wide functional connectivity maps of spontaneous activity measured by seed-based rs-fMRI and fMRI with cortical inactivation (with Fig. 1A Opto paradigm). Brain atlas drawings (leftmost images) show seed ROIs and optogenetic stimulation sites. **(B)** Strong bilateral homotopic correlations were observed in seed-based rs-fMRI for the S1FL, M1 and S2 (wild-type, n=5) and in seed-based rs-fMRI of transgenic VGAT-ChR2 mice (Fig. S4A). However, **(C)** ipsilateral somatosensory regions, including the cortices and thalamic nuclei, mostly responded to optogenetic cortical inactivation of the S1FL (VGAT-ChR2, n=8), M1 (n=7) and S2 (n=7), consistent with neuronal projections (Fig. S4B). **(D-F)** Connectivity matrices of the somatosensory network measured by rs-fMRI vs. fMRI with cortical inactivation vs. neuronal tracing. For the relative connectivity strengths in somatosensory-related ROIs in each hemisphere, **(D)** degrees of resting-state connectivity were normalized to the strongest connectivity with seed areas, **(E)** fMRI signal changes were normalized to those in the optogenetic source region, and **(F)** anatomic projection densities were normalized to those in injection site (obtained from the Allen Institute; (19)). Resting-state fMRI based on temporal correlation mainly reflected bilateral homotopic connectivity, whereas spontaneous connectivity measured by fMRI with cortical inactivation and anatomical connectivity were mainly observed in ipsilaterally networked cortical and subcortical regions, indicating that the bilateral homotopic connectivity of rs-fMRI was unlikely to be due to direct corticocortical neural communication. Red-to-orange color scale, relative connectivity strength with the seed or source areas.

**Figure 3.**
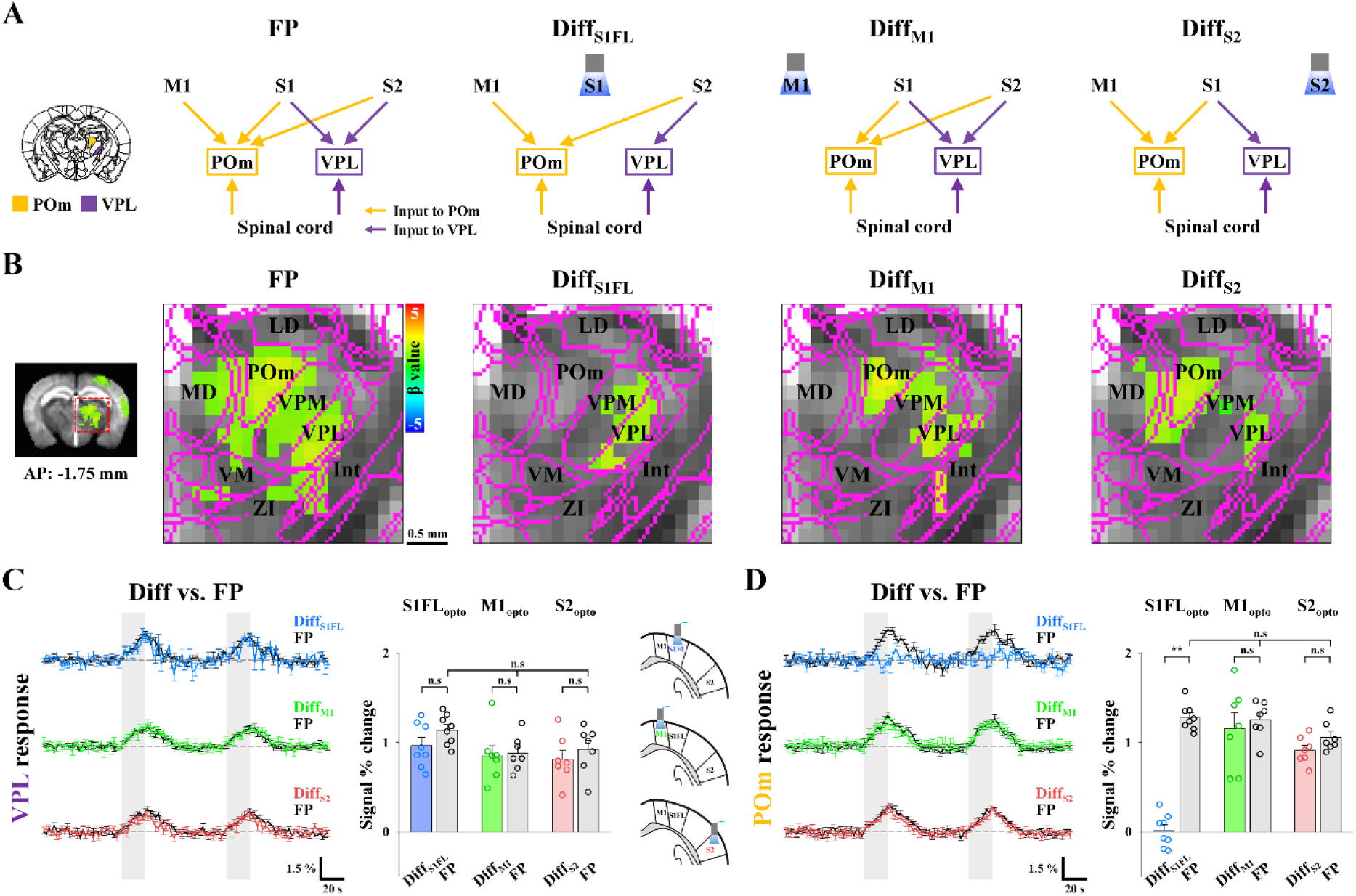
Dissection of somatosensory-driven thalamic fMRI responses with cortical silencing. **(A)** Schematic diagrams to determine the contribution of long-range inputs to somatosensory-evoked thalamic fMRI responses by means of optogenetic cortical inhibition. Functional activities in thalamic nuclei during forepaw stimulation (FP) can originate from multiple long-range inputs via ST and CT pathways, whereas differences in thalamic fMRI responses (Diff with a subscript indicating the inactivation site) between optogenetic cortical inactivation with and without sensory stimulation are related to all possible inputs except those from the target cortex of the S1FL, M1 and S2 to the thalamic nuclei; thus, with comparison of those conditions (Diff vs. FP), the contribution of each pathway to thalamic responses was investigated. VPL, ventral posterolateral nucleus; POm, posterior medial nucleus. **(B)** fMRI maps of the thalamic nuclei. Forepaw stimulation (FP) induced activity in two distinct foci, VPL and POm (FP: n=22). Somatosensory-evoked activities in the VPL were maintained in the fMRI difference maps (Diff) between optogenetic cortical inactivation with and without FP, whereas POm activity disappeared in the difference map of S1FL inactivation (Diff_S1FL_: n=8) but not M1 (Diff_M1_) and S2 (Diff_S2_) inactivation (n=7, each). LD, lateral dorsal nucleus; MD, mediodorsal nucleus; POm, posterior medial nucleus; VPM, ventral posteromedial nucleus; VPL, ventral posterolateral nucleus; VM, ventral medial nucleus; Int, internal capsule; ZI, zona incerta; scale bar, 0.5 mm. **(C-D)** Contribution of functional pathways to somatosensory-evoked VPL and POm responses. fMRI time courses in **(C)** the VPL and **(D)** the POm for the difference between combined stimulation and cortical silencing (Diff_S1FL_, blue lines; Diff_M1_, green lines; Diff_S2_, red lines) and those for FP only (FP, black lines) were compared. **(C)** Under cortical silencing with and without FP, those differential responses (colored bar) are similar to the total VPL responses (gray bar), **(D)** whereas the sensory-evoked POm response was completely suppressed during S1FL silencing (blue bar) but not during M1 (green bar) and S2 (red bar) silencing. Therefore, the VPL has dominant inputs from the spinal cord, not the cortex, and POm activity is directly driven by the S1FL, not by the spinal cord. Gray vertical bar in time courses, 20-s stimulus; error bars, SEM; **p < 0.01 (paired t-test).

**Figure 4.**
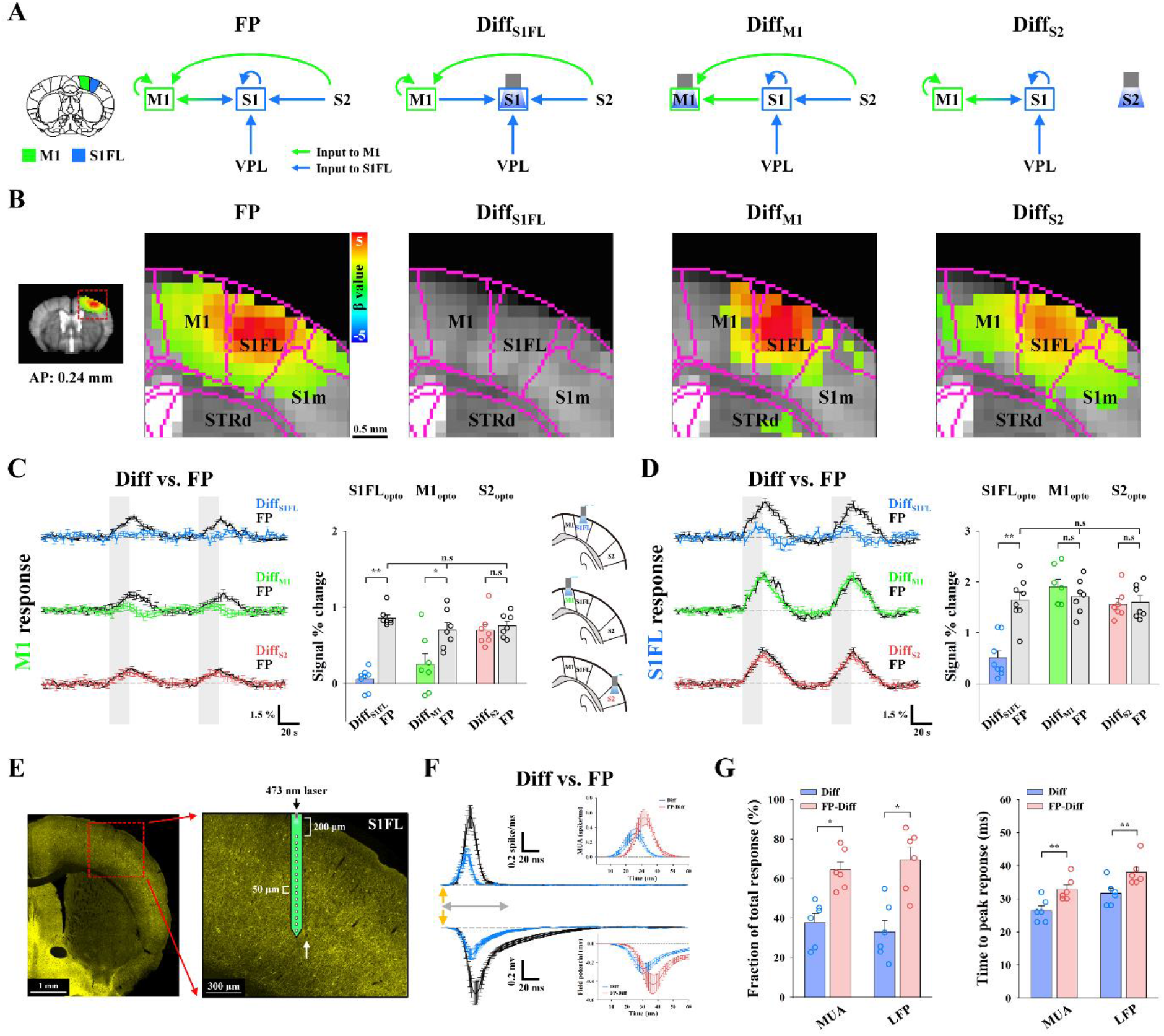
Dissection of TC, CC and IC activity during somatosensory fMRI responses in the S1FL and M1. **(A)** Schematic diagrams to determine the contribution of long-range and local activity to somatosensory-evoked M1 and S1FL responses with optogenetic cortical inhibition. Functional activity in the S1FL during forepaw stimulation (FP) can originate from both long-range inputs via TC and CC pathways and local IC activity, whereas M1 responses can comprise CC inputs and IC activity. The differences in cortical responses between optogenetic cortical inactivation with and without sensory stimulation (Diff) are related to input-driven activity, excluding CC and/or IC activity from the inactivation site (indicated as a subscript). M1, primary motor area; S1FL, primary somatosensory area of the forelimb. **(B)** fMRI maps in cortical areas including M1 and the S1FL. Somatosensory-evoked activities in M1 (FP: n=22) disappeared in the fMRI difference map between M1 (Diff_M1_: n=7) or the S1FL (Diff_S1FL_: n=8) optogenetic inactivation with and without forepaw stimulation but remained after S2 inactivation (Diff_S2_: n=7), whereas somatosensory-evoked S1FL responses disappeared only after S1FL inactivation. M1, primary motor area; S1FL, primary somatosensory area of the forelimb; S1m, primary somatosensory area of mouth; STRd, striatum of dorsal region. **(C-D)** Contribution of functional pathways to somatosensory-evoked M1 and S1FL responses. fMRI time courses in **(C)** M1 and **(D)** S1FL to determine the difference between combined stimulation and cortical silencing (Diff_S1FL_, blue lines; Diff_M1_, green lines; Diff_S2_, red lines) and those for forepaw stimulation (FP, black lines) only were compared. **(C)** The differential response in M1 (green bar) between M1 silencing with and without forepaw stimulation was approximately one-third of the total M1 response (gray bar). In addition, the sensory-evoked M1 response was mostly suppressed after S1FL silencing (blue bar) but not after S2 silencing (red bar). **(D)** The difference in S1FL responses between S1FL silencing with and without forepaw stimulation (blue bar) is related to TC input, which accounts for approximately one-third of the total S1FL response (gray bar). However, sensory-evoked S1FL responses were not changed by M1 (green bar) and S2 (red bar) silencing. Therefore, M1 fMRI response were dominantly driven by the S1FL, and S1FL fMRI responses were directly driven by the VPL, with minimal corticocortical contribution; both of these responses were amplified by intracortical circuits. Gray vertical bar in time courses, 20-s stimulus; error bars, SEM; *p < 0.05 and **p < 0.01 (paired t-test). **(E-G)** Electrophysiological recording at 200–1000 µm depth with the 16-channel opto-electrode performed to investigate the neural source of the S1FL fMRI response. **(E)** Fluorescence microscopy image from a representative animal shows the expression of ChR2-EYFP (yellow) in the S1FL and the location of the electrode track (white arrow). **(F)** Neural data were time-lock averaged for the forepaw stimulus interval of 250 ms. Depth-averaged neural responses (MUA and LFP, VGAT-ChR2, n=6) in the S1FL for the difference between combined stimulation and S1FL silencing (Diff, blue lines) and those for forepaw stimulation (FP, black lines) only were compared to isolate the local IC activity (FP-Diff, red lines in inset). Detailed depth-dependent responses are shown in *SI Appendix*, Fig. S8. **(G)** Fraction and time to peak of sensory inputs (blue bar) and IC activity (red bar) in MUA and LFP. Quantitative MUA and LFP values are reported in Table S2 and S3. Yellow arrows in **(F)**, 0.5-ms forepaw stimulus; gray double arrows in **(F)**, 60-ms period for the inset figure; error bars, SEM; *p < 0.05 and **p < 0.01 (paired t-test).

## Results

To map brain-wide spontaneous and evoked long-range somatosensory networks by fMRI, VGAT-ChR2-EYFP mice expressing light-sensitive opsin proteins in GABAergic interneuron populations were used (Fig. S1 for expression in inhibitory neurons). An optical fiber cannula (105-µm inner core diameter) was implanted into the middle of the cortex in the S1FL or S2 and the upper cortical area in M1 (Fig. S2A) to silence local pyramidal neuronal activities via optogenetic stimulation of inhibitory neurons with blue light (473 nm, 3 mW, 10 ms duration, and 20 Hz) (Fig. 1A).

The suppression of neural activity was confirmed by recording MUAs with 16-channel optoelectrode under the same photostimulation conditions used for fMRI studies. During 20 s of photostimulation to the cortical surface of the S1FL, an increase in stimulation-induced inhibitory neuronal activity mostly suppressed spontaneous activity across all cortical depths (Fig. 1B; baseline (black profile and bar) vs. light on (blue profile and bar); n=6 mice), which is consistent with previous findings (33, 49). Similar uniform suppression was observed across all cortical depths when photostimulation was targeted at a depth of 500 µm in the S1FL (Fig. S2B). With 3 mW optogenetic stimulation of VGAT-ChR2, the spatial spread of inhibition was expected to be >1 mm from the center of fiber (49), which covers the S1FL, M1, or S2 areas.

Optogenetic fMRI of anesthetized VGAT-ChR2 mice was performed with an ultrahigh magnetic field of 15.2 T. Fiber implantation often causes image distortions and signal dropouts in EPI scans commonly used for fMRI studies, which are more severe with higher magnetic fields (see red box of Fig. 1C; Fig. S3A). Thus, we adopted a conventional gradient-echo imaging technique with a short TE of 3 ms to minimize TE-sensitive BOLD contributions and image distortions. This approach allowed us to identify fiber positions (see blue box of Fig. 1C; Fig. S3A) and coregister brain MRI scans with the mouse brain atlas. To enhance functional sensitivity and specificity, 45 mg/kg superparamagnetic iron oxide nanoparticles were injected into the blood of the animals (Fig. S3B for selection of the iron oxide dose), which resulted in CBV sensitization. An increase in CBV increased the amount of iron oxide within the voxel, thus decreasing fMRI signals. Therefore, the change in the polarity of the original CBV-weighted fMRI signal was inverted to match that of the CBV responses. Multislice CBV-weighted fMRI scans were obtained with spatial resolution = 156 × 156 × 500 μm^3^ and temporal resolution = 2 s for mapping functional connectivity with highly specific blood volume responses. Photostimulation did not induce heating-related fMRI artifacts in naïve mice (Fig. 1D; C57BL/6J, n=3 mice); thus, fMRI responses to optogenetic stimulation genuinely reflected underlying neural activity.

To dissect the functional circuits by fMRI, optogenetic cortical inhibition was adopted during somatosensory stimulation (Fig. 1E). Silencing cortical excitatory neurons by optogenetically activating ChR2-expressing inhibitory neurons suppresses local excitatory recurrent circuits and outputs to downstream pathways (15, 50) (Opto in Fig. 1E), resulting in a decreased fMRI signal in local and downstream networked areas (deactivation). Notably, most cortical GABAergic neurons do not project other brain regions. Therefore, this approach can be used to map spontaneous neural interactions at rest. Simultaneous forepaw stimulation during cortical silencing can preserve sensory-evoked inputs to the thalamus and cortex via upstream spinothalamic (ST) and TC pathways with downstream suppression (Comb in Fig. 1E). Since inhibitory neural activation induces a hemodynamic response at the stimulation site (41, 51) and in downstream regions, the contribution by this type of activation should be removed. It is assumed that the common photostimulation-driven hemodynamic response is completely eliminated by calculating the difference between the optogenetic fMRI responses observed with and without sensory stimulation (Comb – Opto = Diff in Fig. 1E). Therefore, the remaining signal is due to the contribution of upstream ST and TC pathways in the absence of local recurrent circuits and downstream projections from a given target region. Then, the contribution of downstream suppression of fMRI responses can be determined from somatosensory-evoked fMRI responses (Diff vs. forepaw somatosensory stimulation (FP) in Fig. 1E), allowing us to determine the functional causality for long-range and local processing in somatosensory networks. Although a similar circuit-analysis approach of sensory stimulation with and without local silencing has been used for electrophysiological studies in preselected areas (13–15), fMRI with and without optogenetic inhibition can be used to determine brain-wide long-range circuits at a population level.

### Mapping somatosensory networks of resting-state activity: rs-fMRI *vs*. cortical silencing fMRI

First, we investigated resting-state somatosensory networks by fMRI (Fig. 2). Pertinently, rs-fMRI connectivity, which is commonly used, is determined by synchronization of fluctuating fMRI signals between regions in the absence of a task, which is presumably related to the intrinsic network of spontaneous activity. Alternatively, optogenetic cortical silencing (in a block-design paradigm with and without 20-s optogenetic stimulation, Fig. 1A Opto paradigm) suppresses spontaneous output activity from the stimulated site and causally reduces input to downstream networked areas (see actual time courses in Fig. S6B-D). Thus, the decrease in fMRI responses due to cortical silencing is closely related to the strength of resting-state connectivity between the optogenetic stimulation site and the connected regions.

Rs-fMRI data for 10-min scans (i.e., 300 volumes) without stimulation were obtained from five naïve mice; in addition, rs-fMRI connectivities in the S1FL, M1, and S2 seed regions of interest (ROIs) in the right hemisphere were mapped (Fig. 2B). Commonly observed strong bilateral homotopic cortical connectivity was detected without a significant network between the seed ROIs and the ipsilateral thalamus (Fig. 2B). Similar homotopic cortical connectivity was observed in transgenic VGAT-ChR2 mice (see Fig. S4A), indicating that bilateral homotopic correlation is a common feature in rs-fMRI scans. This bilateral cortical connectivity may have been due to direct corticocortical communications and/or synchronized common neural and vascular sources. Therefore, fMRI with cortical inactivation can be used to examine the contribution of spontaneous neuronal CC communications to bilateral rs-fMRI connectivity.

Focal silencing of S1FL, M1, or S2 activity by 20 s of optogenetic stimulation of inhibitory neurons reduced CBV in networked cortical and subcortical sites (Fig. 2C; Fig. S6). Inactivation of the S1FL induced negative CBV changes at the ipsilateral S1 (hindlimb and whisker barrel); M1; S2; thalamic nuclei, including VPL and POm; and striatum (see Fig. 2C for group-averaged fMRI maps; Fig. S6B for responses of individual animals; VGAT-ChR2, n=8 mice). Similar observations were detected for M1 inactivation (see Fig. 2C for group-averaged fMRI maps; Fig. S6C for responses of individual animals, n=7 mice) and for the inhibition of S2 activity (Fig. 2C for group-averaged fMRI maps; Fig. S6D for responses of individual animals, n=7 mice). Interestingly, the S1FL and M1 had reciprocal connections with similar magnitudes, while the CT connection was stronger in the POm than in the VPL (Fig. 2C). These fMRI networks are topologically consistent with anatomical neural tracing studies (see Fig. S4B) (19).

To examine which resting-state somatosensory networks by rs-fMRI (measured by correlation analysis) and cortical silencing fMRI (measured by the signal reduction due to cortical inhibition in a resting state) closely reflect intrinsic neural networks (measured by viral tracer injections), the relative connectivity strengths were quantified by normalization with the strongest connectivity in rs-fMRI (Fig. 2D), with the response at the stimulation site during cortical silencing fMRI (Fig. 2E) and with the projection density in tracer injection site (Fig. 2F). The commonly observed bilateral homotopic connections of rs-fMRI (52, 53) were weakly observed in networks during cortical inhibition fMRI and neural tracing studies. In the degree of correlation with connectivity strength of neural tracing data, rs-fMRI connectivity showed a poor similarity (r=0.09), whereas networks by cortical silencing fMRI were well-matched (r=0.71). These indicate that the dominant bilateral cortical connectivity detected by rs-fMRI was unlikely due to direct neuron-based CC communication. Overall, brain-wide spontaneous networks in the resting state can be mapped by fMRI with inhibition at focal sites.

### Dissection of somatosensory activity-driven thalamic responses by cortical silencing

Somatosensory stimulation of the forepaw (0.5 mA, 0.5 ms duration, and 4 Hz) induced robust CBV increases in the contralateral somatosensory network (see Fig. S5 for group-averaged fMRI maps; Fig. S6A for responses of individual animals, n=22 mice). Brain-wide activation sites were observed in the cortex, including in the S1FL; M1; S2; and thalamic nuclei, including VPL and POm (Fig. S5B). This result suggests that evoked responses to forepaw stimulation possibly contain ST, TC, CC and CT pathways. To dissect the functional circuits that evoked an fMRI response, optogenetic cortical inhibition was performed at the S1FL, M1 and S2 in conjunction with forepaw stimulation.

Initially, we investigated the relative contribution of ST and CT circuits to thalamic fMRI responses (Fig. 3). To separate these two contributions to thalamic fMRI signals, fMRI responses to optogenetic silencing of the targeted cortex (see individual animal data in Fig. S6) were compared to those of simultaneous silencing and forepaw stimulation (see individual animal data in Fig. S7). These differences are directly related to all inputs except the CT inputs from the target site to the thalamic nuclei (Diff_S1FL_, Diff_M1_, and Diff_S2_ in Fig. 3A). In the VPL, the CBV response to somatosensory stimulation was similar to the difference between cortical silencing outcomes and silencing/forepaw stimulation outcomes (Diff), regardless of S1FL, M1, or S2 silencing (Fig. 3B and 3C); this finding suggests that the VPL is driven by inputs from the spinal cord, not from the cortex. In the POm, S1FL silencing suppressed fMRI responses to forepaw stimulation completely, whereas M1 and S2 silencing had no impact (Fig. 3B and 3D), indicating that POm activity was directly driven by S1, not by ST inputs. With our fMRI approach, the relative contributions of feedback CT and feedforward ST activity to sensory-evoked thalamic responses were successfully determined.

### Dissection of somatosensory-evoked S1FL and M1 responses to TC, CC and IC activity

Next, we examined the relative contribution of long-range inputs and local circuits to fMRI responses in the S1FL and M1 (Fig. 4). Somatosensory-induced fMRI responses of the S1FL can contain TC input from the VPL, IC activity and CC feedback from M1 and S2, whereas fMRI responses of M1 can contain CC inputs and IC activity. The role of each pathway in somatosensory-evoked fMRI responses in S1FL and M1 was examined by optogenetic silencing of the S1FL, M1, or S2 (Fig. 4A for input circuit diagrams and Fig. 4B for corresponding fMRI maps; see individual animal traces in Figs. S6 and S7).

We first examined the contributions of functional pathways to somatosensory-evoked M1 responses (Fig. 4C). M1 responses were completely suppressed when optogenetic S1FL silencing was performed (Diff (blue) vs. FP (black/gray) in Fig. 4C) but remained the same when S2 activity was silenced (Diff (red) vs. FP (black/gray) in Fig. 4C). This suggests that somatosensory-evoked M1 activity is mostly driven by CC projections from the S1FL. CC input-driven magnitude responses during M1 silencing accounted for approximately 36% of the total somatosensory-evoked fMRI response in M1 (Diff (green bar) vs. FP (gray bar) in Fig. 4C). Since the input-driven CBV response duration (green line) was shorter than the duration of the total CBV response (black line), the AUCs for these measurements were also compared; this analysis revealed that the input-driven CBV response accounted for 24% of the total response. M1 somatosensory-evoked activity was shown to be driven by the S1FL feedforward projection and amplified via the IC circuitry.

Then, the contributions of the TC, IC and CC circuits to S1FL responses were separated (Fig. 4D). Based on fMRI data acquired after M1 or S2 silencing (green or red), we found that CC projections from M1 and S2 to the S1FL were negligible during forepaw stimulation. This result indicated that there was no feedback CC long-range contribution to the sensory response in the S1FL, as expected under anesthesia (54, 55). When IC activity in the S1FL was suppressed, a positive CBV response remained, with a reduced magnitude of approximately 31% of that of the total stimulus-driven response (Diff (blue) vs. FP (black/gray) in Fig. 4D) and 26% of that of the total AUC.

To investigate the neural source of the somatosensory-induced fMRI reduction caused by local silencing, 16-channel electrophysiological recordings with the same stimulus paradigms as fMRI (Fig. 4E, VGAT-ChR2, n=6 mice) were acquired in the S1FL. Time-locked neural data for one forepaw stimulus pulse interval of 250 ms were obtained by averaging all 250-ms windows (FP; black in Fig. 4F). Inhibitory neuronal activities induced by optogenetic stimulation of GABAergic neurons were subtracted from the difference between those responding to optogenetic stimulation with and without somatosensory stimulation (Diff; blue). MUAs and LFPs were compared across different experimental conditions (see Fig. 4F and 4G for averaged responses across all 16 channels and Fig. S8 for depth-dependent responses). The difference between optogenetic silencing of S1FL and simultaneous silencing/forepaw stimulation (Diff; blue) indicated neural activities induced by TC input, whereas somatosensory-driven activity (FP) contained both long-range input and local recurrent activity (Fig. 4F). Thus, the local recurrent activity (FP-Diff; red) can be obtained from the difference between long-range TC input (Diff; blue) and total somatosensory activity induced by forepaw stimulation (FP; see inset figures in Fig. 4F). The thalamic input activity integrated over time was ∼35% of the forepaw-induced neural activity (Fig. 4G; 37.73 ± 4.54% for MUA and 33.03 ± 5.92% for LFPs; Table S2 for MUA and Table S3 for LFP), which is in good agreement with the fMRI observations (31% for magnitude, 26% for AUC). The somatosensory-evoked local recurrent activity suppressed by optogenetic silencing occurred ∼6 ms later than the input activity (Fig. 4G; Diff vs. FP-Diff; 26.50 ± 1.43 ms vs. 32.83 ± 1.45 ms for MUA; 31.67 ± 1.52 ms vs. 38.00 ± 1.71 ms for LFPs; Table S2 for MUA and Table S3 for LFPs). Combined with macroscopic fMRI and microscopic electrophysiology data, these findings showed that somatosensory-evoked TC activity was amplified ∼2-fold in the S1FL due to local recurrent circuits (15), and demonstrated that fMRI can be used to separate long-range input and local circuit contributions.

Then, we examined whether TC and CC input layers in the S1FL and M1 can be identified by upsampled cortical depth-dependent CBV responses (Fig. 5). Somatosensory-evoked CBV responses peaked at layer 4 (L4) of the S1FL (Fig. 5B and 5C for group-averaged fMRI responses; Fig. S9A-i for responses of individual animals, n=22 mice), indicating the TC input, and peaked at layer 2/3 (L2/3) of M1 (Fig. 5B and 5D for group-averaged fMRI responses; Fig. S9 for responses of individual animals, n=22 mice), indicating the CC input. Cortical suppression in the S1FL, M1 and S2 induced negative CBV changes in the optogenetic simulation site and peak responses in CC input layers of networked regions (Fig. 5B-D for group-averaged fMRI responses; Fig. S9 for responses of individual animals). When M1 was suppressed, the CC projection occurred at L2/3 and L5A, not L4, in S1 (56). Corresponding CBV peaks with different magnitudes were detected, with a small dip at ∼150 µm-thick L4 (see Fig. 5C green), indicating that CBV responses exhibit a hemodynamic point spread function (PSF) of < 150 µm FWHM. Our laminar-resolved fMRI data showed that cortical CBV responses peaked at synaptic input layers with high specificity.

**Figure 5.**
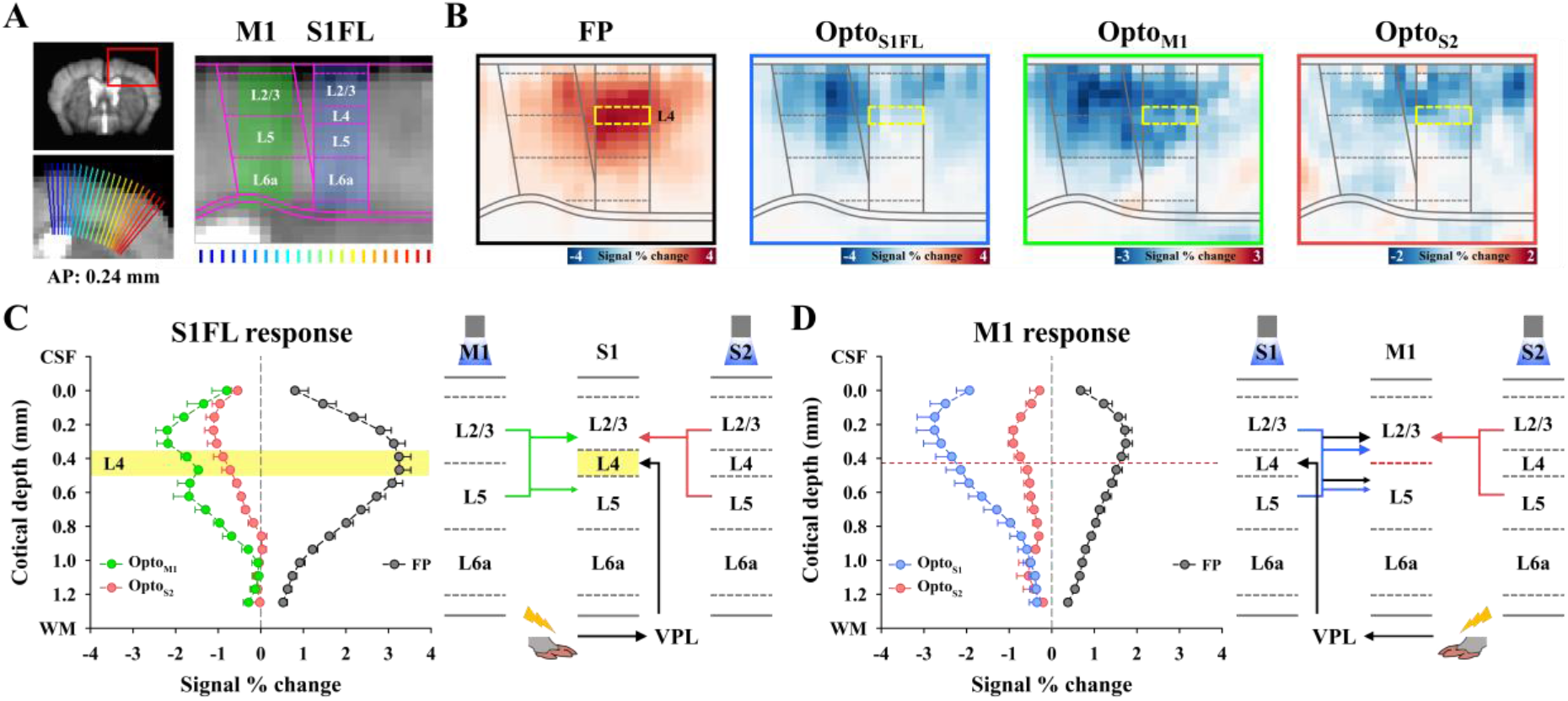
The highest CBV responses at synaptic input layers in the S1FL and M1. **(A)** Cortical flattening for layer-specific fMRI analysis in the S1FL and M1. The cortical area (red box) was upsampled to double using bicubic interpolation and linearized using radially projecting lines (blue to red) perpendicular to the cortical edges (underlay, study-specific brain template; overlay, Allen mouse brain atlas). Laminar boundaries for each cortex were defined as the laminar thickness distribution. **(B)** Cortical depth-dependent fMRI maps with nominal 78 µm in-plane resolution for somatosensory-evoked and spontaneous activities. Somatosensory-evoked activities (FP: n=22) were localized at L4 of the S1FL (yellow-dashed box) and L2/3 of M1. Strong suppression occurred at L2/3 of the S1FL and M1 during optogenetic cortical inactivation (Opto_S1FL_: n=8, Opto_M1_: n=7, Opto_S2_: n=8). **(C-D)** Cortical depth-dependent CBV profiles and expected circuit diagrams. Signal changes were averaged for the same depth and plotted as a function of distance from the surface in **(C)** the S1FL and **(D)** M1. **(C)** The somatosensory-evoked response (black) peaked at L4 of the S1FL, while M1 inactivation (green) induced two cortical peaks at L2/3 and L5; in addition, S2 silencing (red) induced the largest changes at L2/3. Since the ∼150-µm-thick L4 of the S1FL does not receive inputs from M1, the cortical profile responding to M1 stimulation indicates that the PSF of laminar CBV response is less than 150 µm FWHM. **(D)** The largest fMRI changes in M1 were mostly detected at L2/3, regardless of the type of stimulus used. Individual animal profiles are shown in Fig. S9.

### Separation of the TC, CC, and cortico-thalamo-cortical (CTC) contributions to somatosensory-evoked S2 responses

Finally, we examined the relative contribution of long-range TC, CC and CTC inputs and local circuits to fMRI responses in S2 (Fig. 6). The secondary somatosensory area is anatomically connected to the VPL, POm, S1FL, and M1 (57). Thus, functional inputs to S2 during forepaw stimulation can originate from monosynaptic circuits from the VPL (TC) and S1FL (CC) and disynaptic projections from the S1FL via M1 (S1FL → M1 → S2) and the POm (S1FL → POm → S2). When M1 activity was suppressed, fMRI responses at S2 remained the same (Diff (green) vs. FP (black/gray) in Fig. 6B), indicating that the contribution of the S1 → M1 → S2 pathway was negligible. The contribution of the TC projection, calculated by the difference between the signal change without and with S1FL silencing (Figs. S6 and S7 for individual animal traces), constituted 25% of the total S2 response (Diff (blue) vs. FP (black/gray) in Fig. 6B), whereas the remaining 75% of inputs originated from the S1FL. When the sensory-evoked S2 fMRI responses were compared without and with S2 silencing, CBV responses of long-range synaptic inputs accounted for approximately 28% of the total percent change in S2 (Diff (red) vs. FP (black/gray) in Fig. 6B; 30% of the total AUC), whereas the remaining responses were due to local circuit contributions. These findings suggested that somatosensory-evoked S2 activity was mostly driven by feedforward S1FL inputs with minor VPL inputs and amplified by the IC recurrent circuit.

**Figure 6.**
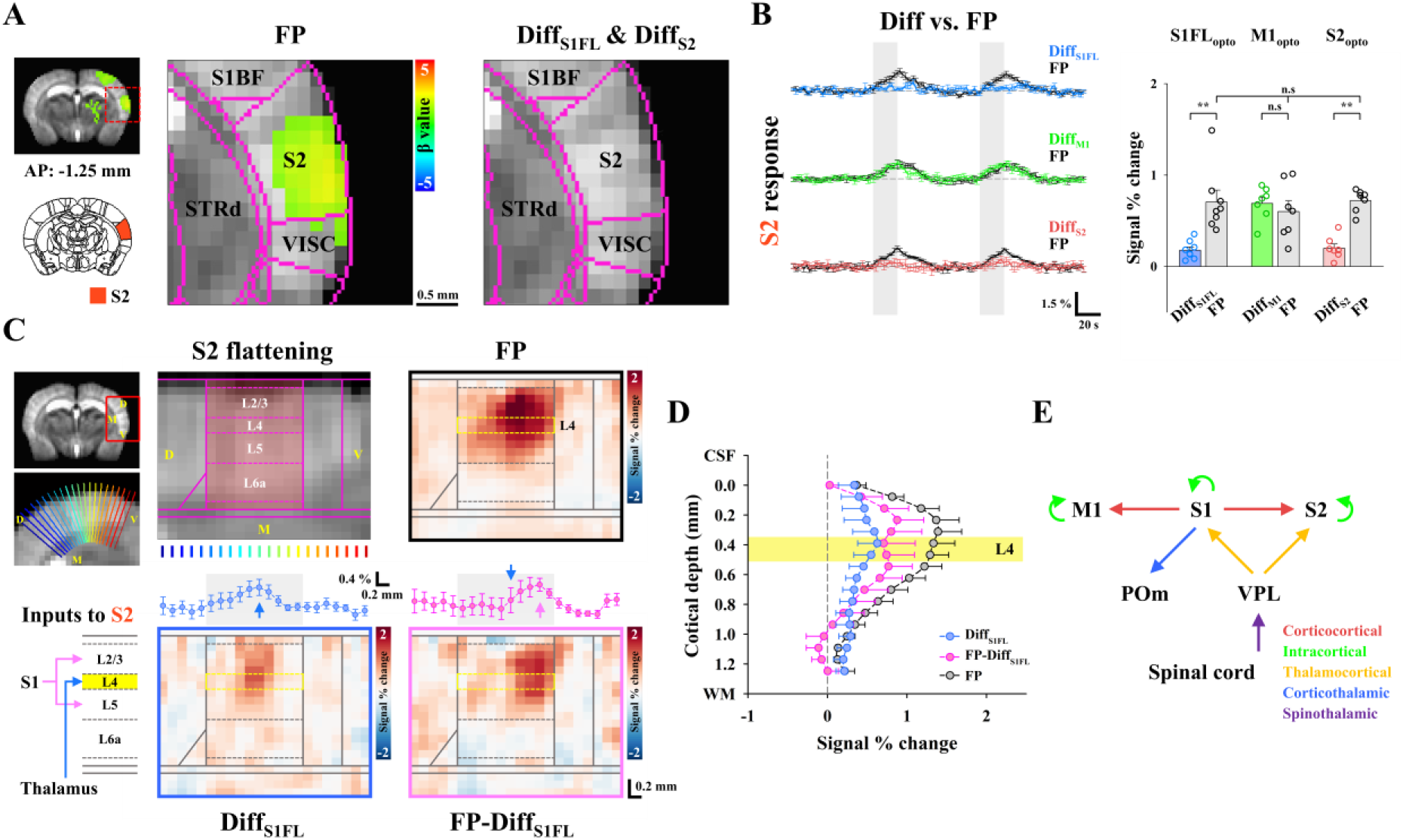
Separation of TC, CC, and CTC contributions to sensory-evoked fMRI responses in S2. **(A)** fMRI maps in cortical areas including S2. Somatosensory-evoked activities in S2 (FP: n=22) disappeared in the fMRI difference map between S1FL or S2 optogenetic inactivation with and without forepaw stimulation (Diff_S1FL_: n=8, Diff_S2_: n=7) but remained after M1 inactivation (not shown here). S1BF, primary somatosensory area of the barrel field; S2, secondary somatosensory area; VISC, visceral area; STRd, striatum of dorsal region. **(B)** Contribution of functional pathways to the somatosensory-evoked S2 response. fMRI time courses in S2 for the difference between combined stimulation and cortical silencing (Diff_S1FL_, blue line; Diff_M1_, green line; Diff_S2_, red line) and those obtained for forepaw stimulation only (FP, black lines) were compared. Under S2 silencing with and without forepaw stimulation, the difference response in S2 (red bar) was approximately one-third of the total S2 response (gray bar). The sensory-evoked S2 response was largely suppressed in S1FL silencing (blue bar) but still had TC input (∼25% of the total S2 response) and was not changed by M1 silencing (green bar). Therefore, the S2 fMRI response was predominantly driven by the S1FL with minor VPL contribution and was amplified by intracortical circuits. Gray vertical bar in time courses, 20-s stimulus; error bars, SEM; **p < 0.01 (paired t-test). **(C)** Layer-specific fMRI analysis in S2. Thalamic inputs, including both TC and cortico-thalamo-cortical (CTC) circuits, project to L4, whereas direct S1FL inputs project to L2/3 and L5 (Inputs to S2). To investigate the laminar origin of the evoked S2 response, the S2 area was upsampled and linearized using radially projecting lines (blue to red) perpendicular to the cortical edges (underlay, study-specific brain template; overlay, Allen mouse brain atlas). Laminar boundaries were defined as cortical thickness distribution (S2 flattening). To separate layer-specific TC inputs and direct/indirect S1FL inputs to S2 from the total sensory-evoked response, fMRI responses for the difference between combined stimulation and S1FL silencing (Diff_S1FL_, TC inputs) and those for forepaw stimulation only (FP from only dataset paired with Opto_S1FL_) were compared (FP-Diff_S1FL_, direct/indirect S1FL inputs). In S2 flattened maps, somatosensory-evoked responses were averaged across the cortical layers and plotted in a dorsal-to-ventral direction (left-to-right). The patches responding to TC (blue profile in Diff_S1FL_) and direct/indirect S1FL (pink profile in FP-Diff_S1FL_) inputs were separable. Blue (Diff_S1FL_) and pink (FP-Diff_S1FL_) arrows, locations of the response peak; error bars, SEM. **(D)** For the cortical depth-dependent profile, signal changes were averaged for the same depth and plotted as a function of distance from the surface in S2. During the sensory-evoked response in S2 (FP, black line), the laminar profile for TC inputs peaked at L4 of S2 (Diff_S1FL_, blue line), while the cortical profile projected from S1FL was observed to have double-peak responses at L2/3 and L5 (FP-Diff_S1FL_, pink line), indicating direct CC inputs. Yellow region, layer 4; error bars, SEM **(E)** Putative functional circuits in the somatosensory network as measured by CBV-weighted fMRI with optogenetic cortical silencing.

The synaptic inputs from the S1FL to S2 can be direct S1FL (CC) inputs and/or indirect S1FL (CTC) inputs via POm (58). Thus, our next question was whether the direct S1FL input to S2 (S1FL → S2) can be separated from CTC input (S1FL → POm → S2). Since direct S1FL inputs project to layers 2/3 (L2/3) and L5 and CTC inputs project to L4 (layer “Inputs-to-S2” schematic in Fig. 6C) (56, 58, 59), upsampled cortical depth-dependent analysis can provide an indication of whether CTC inputs are dominant in S2 fMRI responses (Fig. 6C). Since laminar CBV responses are specific to synaptic input layers with a PSF of < 150 µm, its peak position provides information on cortical input layers. During forepaw somatosensory stimulation, laminar S2 responses were broad around L2/3 and L4 (FP in Fig. 6C and the black profile in Fig. 6D). To obtain somatosensory-evoked fMRI signals originating from the VPL, the difference in cortical responses between S1FL inactivation with and without forepaw stimulation was obtained (Diff_S1FL_ in Fig. 6C and blue profile in Fig. 6D). Then, the S1FL input (FP-Diff_S1FL_ in Fig. 6C and the pink profile in Fig. 6D) to S2 was determined by subtracting the VPL input (Diff_S1FL_) from the total sensory-evoked response (FP). The TC inputs peaked at L4 in the medial S2 (Diff_S1FL_ in Fig. 6C and blue profile in Fig. 6C), whereas the inputs from the S1FL induced double peaks at L2/3 and L5 in the ventral S2 (FP-Diff_S1FL_ in Fig. 6C and the pink profile in Fig. 6D), indicating direct CC projections. These TC and CC inputs to S2 are spatially segregated by approximately 0.47 mm in a dorsal-ventral direction (Diff_S1FL_ (blue arrow) vs. FP-Diff_S1FL_ (pink arrow) in Fig. 6C), which cannot be easily identified by microscopic tools with a small field of view. Based on our laminar-resolved CBV fMRI with cortical silencing, we successfully confirmed TC and direct CC inputs to S2. Overall, we completely scrutinized the relative contributions of somatosensory-driven long-range and local recurrent circuits to fMRI responses with the aid of focal optogenetic silencing and laminar-specific CBV contrasts (Fig. 6E).

## Discussion

We developed fMRI approaches with highly specific CBV contrasts and local optogenetic silencing to dissect resting-state and functional brain-wide long-range networks. Pertinently, we successfully determined the relative contribution of each somatosensory circuit to fMRI responses in mice at the population level. VPL responses originated mostly from the spinal cord, whereas the POm received functional CT projections from the S1FL (60, 61). The S1FL received somatosensory input from the VPL (58, 62), and M1 received feedforward CC input from the S1FL. S2 received mostly direct CC input (∼75%) from the S1FL and a small amount of TC input (∼25%) from the VPL. The long-range synaptic input in cortical areas was amplified approximately 2-fold by local IC circuits. Since our findings are consistent with electrophysiology studies at preselected sites (13-15, 22, 24, 58, 60), the fMRI approach to long-range circuit analysis is viable for mapping long-range functional circuits in the whole brain.

The fMRI approach with cortical silencing can be extended for investigations of brain-wide functional circuits employing external stimuli or direct cortical stimulation. Evoked stimulation can be achieved via external or intracortical approaches with optogenetic or electric stimulation, activating entire networks without knowledge of the exact flow direction except for the target site. Thus, localized silencing is necessary to suppress downstream networks; this suppression can be achieved by optogenetic stimulation with high temporal specificity, and pharmacological and photochemical tools for bulk interventions for specific neurotransmitter inputs to a defined brain region with low temporal specificity (10). The advantage of optogenetic tools is that they provide users with the ability to perform fMRI experiments under external stimulation with and without cortical silencing in an interleaved manner; this ability is essential for signal averaging without bias by slowly modulating animal physiology during imaging studies.

### Focal inhibition by optogenetic stimulation of inhibitory neurons

Optogenetic activation of cortical GABAergic neurons has been widely used in circuit/systems neuroscience (15, 50). To identify the contribution of neural circuits to fMRI responses with optogenetic stimulation of the selected region in VGAT-ChR2 mice, it was assumed that cortical GABAergic interneurons were mostly local. Based on Allen mouse brain connectivity (http://connectivity.brain-map.org/projection/experiment/167441329), long-range projections of GABAergic interneurons were found to be negligible in the S1 barrel field. Our assumption appears to be valid in the somatosensory cortex.

Another assumption was that optogenetic stimulation of GABAergic interneurons fully suppresses excitatory recurrent neuronal activities across cortical depths and downstream projections. Although the penetration depth of blue light is dependent on laser power, it is quite shallow (depth of half maximum=∼300 µm (49)). In our MUA measurements in the S1FL, optogenetic stimulation of inhibitory neurons reduced spontaneous MUA by ∼90% across cortical layers (Fig. 1B and Fig. S2B) and suppressed somatosensory stimulation-induced IC recurrent circuits (Fig. 4F); these outcomes are possibly due to local circuits within a cortical column. In our fMRI studies, the somatosensory-evoked responses in the POm and M1 were completely suppressed when inhibitory neurons in the upstream S1FL were optogenetically stimulated, suggesting that cortical inhibition is effective in blocking downstream projections.

The spatial extent of cortical inhibition is important to determine the partial volume of inhibition within the photostimulated cortical region (S1FL, M1, and S2). Li *et al.* (49) measured the spatial spread of inhibition beyond the target area of the somatosensory cortex in VGAT-ChR2 mice. Neural activity is suppressed by > 70% for 1.5 mW photostimulation and > 90% for 14 mW at 1 mm away from the stimulation site, far beyond the spatial spread of light (0.25-0.5 mm). In our case, with 3 mW optogenetic stimulation of VGAT-ChR2, the spatial spread of inhibition was expected to be >1 mm from the center of the fiber. Since the sizes of S1FL, M1, and S2 in mice are approximately 0.84 × 1.69, 0.82 × 1.47, and 1.56 × 1.84 mm^2^, respectively, based on the Allen mouse brain atlas, the region of interest was less than the area inhibited by optogenetic stimulation of GABAergic neurons. Therefore, our assumption of full inhibition is valid.

### Hemodynamic responses to inhibitory neuronal activity

Hemodynamic fMRI responses are believed to be mostly driven by excitatory activity (8). Although GABAergic interneuron activity is known to regulate local vascular tone by the release of vasoactive mediators (e.g., nitric oxide) (63), their activity also interacts with nearby excitatory neurons in the cortex. When inhibitory neurons were activated in VGAT-ChR2 mice, an increase in CBF and CBV was observed for <5 s of stimulation (41, 64, 65), indicating that inhibitory neurons indeed increase hemodynamic responses. However, the increased inhibitory activity by optogenetic stimulation suppressed excitatory activity (inhibition), which led to a decrease in hemodynamic responses. Therefore, hemodynamic responses may be closely dependent on stimulus frequency and duration. Notably, we recently investigated the contribution of inhibitory neuronal activity to fMRI responses using multimodal measurements with electrophysiology, BOLD fMRI and optical imaging during ChR2 stimulation of inhibitory neurons in VGAT-ChR2 mice (41). Under the same stimulation parameters, biphasic BOLD fMRI and CBV-weighted optical imaging responses were observed at the stimulated site with increased inhibitory and decreased excitatory neuronal activity; an initial small positive change (by increased inhibitory activity) was followed by a prolonged negative response (by suppressed excitatory activity). In our CBV fMRI data, a biphasic response in the S1FL during the stimulation period and poststimulus overshoot were observed. The initial CBV increase was directly due to inhibitory neuron activity, as often seen in hemodynamic studies with short stimulation, and the prolonged negative change was due to the suppression of excitatory neurons (inhibition). A balance between excitation of inhibitory neurons and inhibition of excitatory neurons changes the magnitude and polarity of hemodynamic responses.

### Laminar PSF of CBV responses

The resolving power of laminar fMRI across different layers is closely dependent on fundamental CBV point spread and voxel resolution. Spatial specificity to neuronally active layers is improved with stimulation time up to ∼10 s (66, 67), which may occur due to different dynamics of macro-and microvessels, fast-acting penetrating arterioles and highly specific slow-responding capillaries (68). The CBV PSF was previously measured in the S1FL of mice with CBV-weighted optical imaging (69) and in the rat olfactory bulb with CBV-weighted fMRI (70); the PSF was found to be ∼100 µm FWHM. In our studies with nominal 78-µm in-plane resolution, we can resolve laminar responses within the ∼1 mm-thick somatosensory cortex. Two peaks at L2/3 and L5 in the S1FL were observed during optogenetic stimulation in M1, albeit with interanimal variation (Fig. S9), which was expected based on a previously determined CBV PSF (69).

### Mapping spontaneous neural communication

In rodent rs-fMRI, a strong correlation occurs between bilateral homotopic cortices (52, 53) but not between the cortex and thalamus in the ipsilateral hemisphere. The strong bilateral homotopic correlation is often explained by direct CC connections (52, 71) due to the existence of monosynaptic anatomical projections (19). However, careful evaluation of anatomical tracing data show that ipsilateral projections among the somatosensory networks (including thalamus) are generally larger than contralateral homotopic projections (see Fig. 2F and Fig. S4B) (19). Notably, rs-fMRI fails to detect strong ipsilateral connectivity between the cortex and thalamic nuclei within the somatosensory network; such connectivity was indeed observed in spontaneous connectivity maps generated by focal cortical inhibition. These results suggested that conventional rs-fMRI correlation strength dose not truly reflect anatomical monosynaptic connections, but rather common bilateral fluctuations caused by modulatory cholinergic inputs (72–74), noradrenaline driven by the locus coeruleus (73), and thalamic low frequency (75). Further systematic studies are necessary to determine the origin of rs-fMRI connectivity.

Alternatively, brain-wide spontaneous connectivity can be determined by fMRI with optogenetic silencing. In our studies, ipsilateral connections were predominant among the somatosensory network, while the connection strength between bilateral homotopic regions was approximately 30% of the ipsilateral connection strength (see Fig. 2E). These spontaneous connectivity findings were consistent with anatomical tracing data (see Fig. 2F) (19). Furthermore, ipsilateral somatosensory-related regions responding to cortical inhibition overlapped with sites that were active during somatosensory stimulation. These results indicated that fMRI with cortical silencing can detect functionally networked regions. Spontaneous CT communication to the higher-order thalamic nucleus POm was stronger than that to the relay thalamic nucleus VPL, which was consistent with electrophysiological findings (12, 60).

Spontaneous downstream neural network strength can be measured by fMRI with cortical silencing. In our studies, the spontaneous network strength in S1FL → M1 and S2 (−2.75% and −1.98% of the peak response in L2/3) was larger than the somatosensory-evoked response (1.74% and 1.39% of the peak response). The relatively high spontaneous activity was surprising but may have been due to the use of ketamine for anesthesia (see the subsection of “Effect of anesthesia on resting-state and evoked fMRI”).

The important implication of mapping spontaneous neural communication is the identification of potential circuits that modulate behavior. Optogenetic inhibition has been used to elucidate the involvement of brain regions and specific cell populations associated with behavior. However, the neural circuits involved in behaviors could act through local or downstream circuits. Although anesthesia changes the strength of long-range connections, downstream circuits associated with behavior can be mapped by fMRI.

### Long-range input *vs*. local circuit contributions to sensory-evoked fMRI

Somatosensory-evoked long-range circuits were successfully dissected by fMRI. The CC circuit of S1 and M1 has been extensively investigated anatomically and physiologically (56, 76–78). S1 and M1 are reciprocally connected; neural excitation in S1 rapidly propagates into neurons in L2/3 and L5A in M1, which reciprocally activate neurons in L2/3 and L5A in S1 via a feedback loop (77). In our studies (Fig. 5 for the S1FL and M1 fMRI responses to forepaw stimulation and cortical inhibition), feedforward corticocortical projection from S1 to L2/3 in M1 was observed for evoked and spontaneous conditions; however, reciprocal feedback projection from M1 to L2/3 and L5 in S1 was observed only for spontaneous, not evoked, conditions. It should be noted that underlying neuronal features of the downstream targets in corticocortical projections cannot be determined with our fMRI approach, since synaptic inputs to both excitatory and inhibitory neurons can increase fMRI responses. Similarly, the sensory-evoked feedforward CC projection from S1 to L2/3 and L5 in S2 was observed without the reciprocal feedback projection. Although the S2 contributions to sensory-evoked responses in S1 and M1 were negligible in our studies, these projections may significantly contribute in other behavioral contexts, such as perception and decision-making (79–81). The differences between our study and physiological studies were likely due to anesthesia (54). To examine functional brain-wide circuits relevant to behavior, it is essential to perform fMRI on mice while they are active.

In our studies, the long-range inputs and IC circuit contributions to somatosensory-evoked fMRI responses were separated. The long-range synaptic input-driven fMRI response in the S1FL, S2, and M1 accounted for ∼30% of the total somatosensory-evoked response. Similarly, somatosensory-evoked MUA and LFP in the S1FL were reduced to ∼35% of the total evoked neural activity by S1FL silencing. Therefore, the suppressed response, ∼70% of the total fMRI response (∼65% for MUA and LFP), was related to local neural activity. Our finding of a > 2-fold increase in the IC fMRI signal was highly consistent with electrophysiology data showing an approximately 2.2-fold amplification of IC in the barrel cortex (13), a 2.4-fold amplification of IC in the primary auditory cortex (14) and a 3-fold amplification of IC in the primary visual cortex (15), in which in which TC and IC activities were separated with recordings of excitatory cells in L4 after optogenetic excitation of inhibitory neurons. Overall, fMRI with and without cortical inhibition can be used to identify the contributions of long-range input vs. IC circuits, facilitating the detection of cortical excitatory/inhibitory imbalance and consequent dysfunction of IC recurrent circuits.

We investigated the contribution of long-range cortical inputs to the POm and the VPL in the thalamus, which is consistent with a previous electrophysiology study in which somatosensory-evoked responses in the higher-order POm were drastically reduced after S1 inhibition, while first-order ventral posteromedial nucleus responses were not modulated (60). According to our data, the medial thalamic area adjacent to the POm was active during somatosensory stimulation and S1FL silencing. This observation may have been due to the spillover of POm activity with limited spatial resolution and group averaging and/or due to real activation in intralaminar nuclei, which are associated with multimodal sensory activity (82). Thus, fMRI experiments with higher spatial resolution are needed to determine whether medial thalamic activity is genuine.

### Effect of anesthesia on resting-state and evoked fMRI

Anesthesia affects fMRI responses in both resting and functional states. Pertinently, ketamine is known as a dissociative anesthetic that antagonizes N-methyl-D-aspartate (NMDA) receptors preferentially binding to GABAergic interneurons at a low dose (83, 84). Ketamine disinhibits basal firing of excitatory pyramidal neurons, leading to an increase in electroencephalographic activity, metabolic rate, and cerebral blood flow (85). Ketamine anesthesia induces higher bilateral homotopic calcium-based connectivity with less spatial focality than that observed in the awake condition (86). In the functional state, ketamine (with xylazine) increases recurrent intracortical excitation (87) and cortical fMRI responses (88), possibly due to disinhibition compared to that demonstrated in the awake condition. Overall, it is likely that ketamine anesthesia enhances resting-state and evoked fMRI responses in cortical areas compared to the fMRI responses noted in the awake condition.

## Conclusion

Here, we successfully dissected the brain-wide somatosensory circuits underlying fMRI responses. Our fMRI approach combining stimulation and focal inhibition provides a new avenue for investigating population-based neural circuits throughout the whole brain, allowing longitudinal investigations of functional reorganization caused by neuropathological modifications and learning in individual animals. Circuit-level analysis of whole-brain fMRI will complement conventional microscopic functional circuit studies by elucidating brain-wide, population-based information processing.

## Acknowledgments

This project was funded by the Institute for Basic Science in Korea (IBS-R015-D1). We thank Prof. Joonyeol Lee and Kamil Uludag for critically reviewing an early version of this manuscript, Mr. Andrew You for helpful discussions, and Dr. Hyun-Kyung Lim for confocal imaging of brain slices.

