## Supporting Information for "Dissection of brain-wide spontaneous and functional somatosensory circuits by fMRI with optogenetic silencing"

***Corresponding author:**

Phone number: +82-31-299-4350

**This file includes:**

Figures S1 to S9

Tables S1 to S3

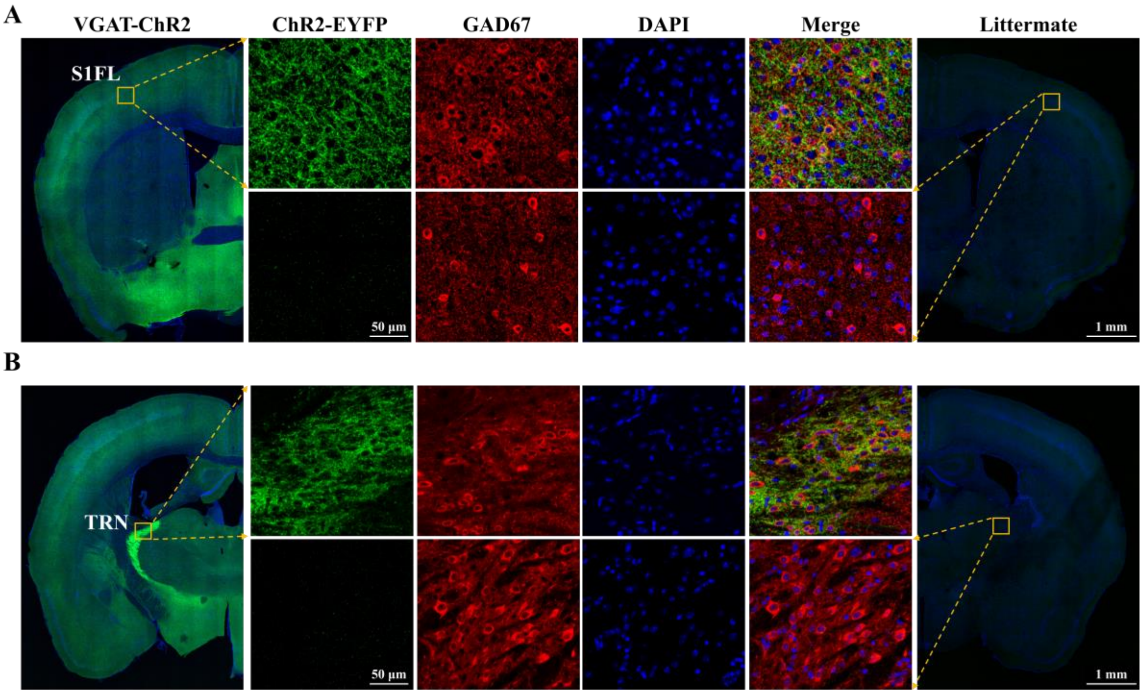


**Figure S1. GABAergic interneuron-specific expression of ChR2-EYFP in VGAT-ChR2 transgenic mouse.**

**(A-B)** Transgenic-specific expression of ChR2-EYFP in **(A)** the primary somatosensory cortex and **(B)** the GABAergic neuron-rich thalamic reticular nucleus of a VGAT-ChR2 positive mouse (the leftmost column and top panel) vs. that of a VGAT-ChR2 negative littermate mouse (the rightmost column and bottom panel). One S1FL-containing slice was chosen to validate the expression of ChR2 in cortical GABAergic neurons. Colocalization of ChR2-EYFP (green) and interneuron markers (GAD67, red) indicates that ChR2 is specific to GABAergic neurons. DAPI was used to visualize cell nuclei (blue). Our histology data are consistent with the reported expression of this transgenic mouse line.

Green-, blue-, and red-colored representation of i, ChR2-EYFP, DAPI and DiI fluorescence, respectively.

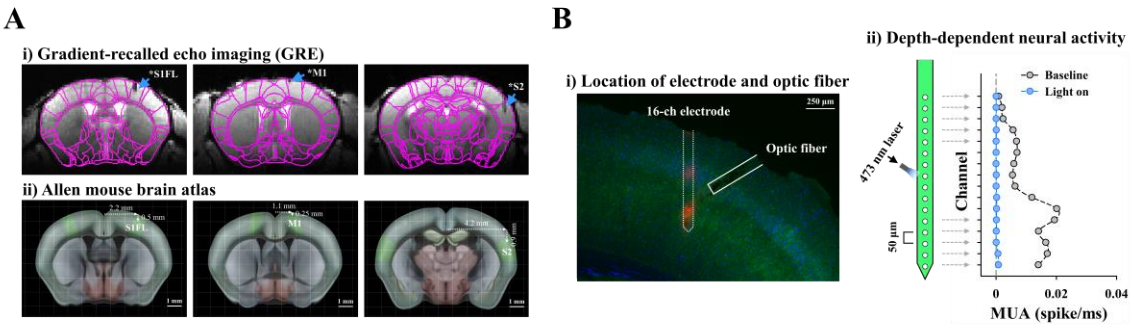


**Figure S2. Fiber position and depth-dependent neural activity in the S1FL**

**(A)** High-resolution mouse brain MR images with fiber-optic implants in the S1FL, M1, and S2 and the corresponding Allen brain atlas. The fiber tip (dark line indicated by a blue arrow) was inserted into the middle of the cortex in the S1FL (0.5 mm from the cortical surface) and S2 and the upper cortical area in M1 (0.25 mm from the cortical surface).

(B) Depth-dependent electrophysiological data in the S1FL without (baseline, gray profile) and with optogenetic stimulation (light on, blue profile) of VGAT-ChR2 transgenic mouse. The 16-channel recording electrode with 50 μm spacing was inserted 1 mm deep, covering a depth between 200 and 1000 μm (dashed line in i with two red spots for electrode position marking), while a 105 μm-diameter optic fiber was obliquely targeted an area 500 μm below the cortical surface (white lined bar in i) to mimic optogenetic fMRI studies in the S1FL. Blue light directly stimulated ChR2-expressing interneurons in the middle and lower cortical layers. However, the MUA was suppressed completely across entire cortical depths, including the upper cortical area above the fiber tip (ii; see also Fig. 1B with cortical surface illumination). This suggests that focal cortical inhibition is effective across all cortical depths, possibly due to recurrent excitatory circuits within a column.

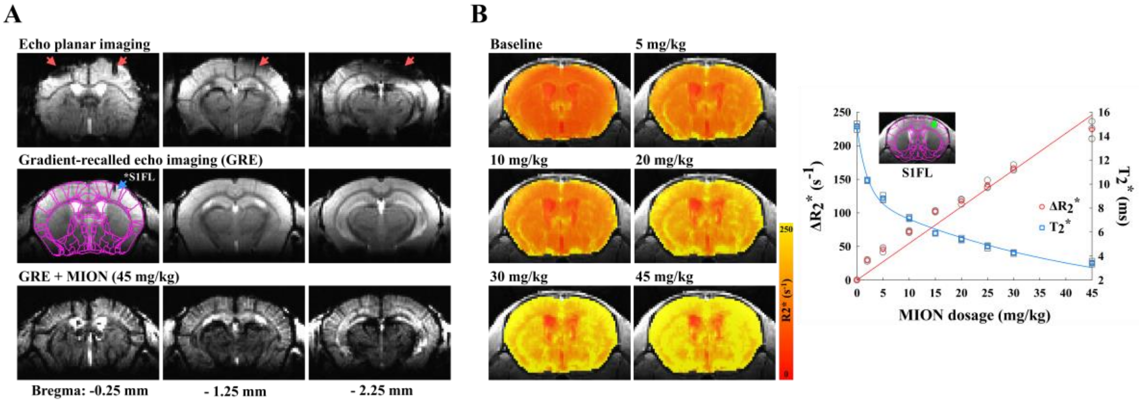


**Figure S3. Imaging quality control for optogenetic CBV-weighted fMRI.**

**(A)** High-resolution mouse brain MR images from one mouse with a fiber-optic implant. Magnetic susceptibility distortion and signal dropout (red arrows) in single-shot gradient-echo EPI images were minimized by adoption of gradient-recalled echo imaging with a short echo time of 3 ms (blue arrow, fiber position). To enhance functional sensitivity and specificity, a 45 mg/kg dose of the superparamagnetic monocrystalline iron oxide nanoparticle (MION) agent was injected into the animals' blood.

**(B)** Contrast agent dose-dependent transverse relaxation rate $R_2^*$ $(= 1/T_2^*)$ changes. The $R_2^*$ maps using multiple gradient-echo imaging (left) were acquired before and after cumulative doses ranging from 2 mg/kg to 45 mg/kg MION in the same animals (wild-type, n=3) to determine the optimal echo time matching to tissue $T_2^*$ for CBV-weighted fMRI. Tissue $T_2$* (blue symbols) and $\Delta R_2^*$ (red symbols) were calculated from S1FL ROI at each MION dose.

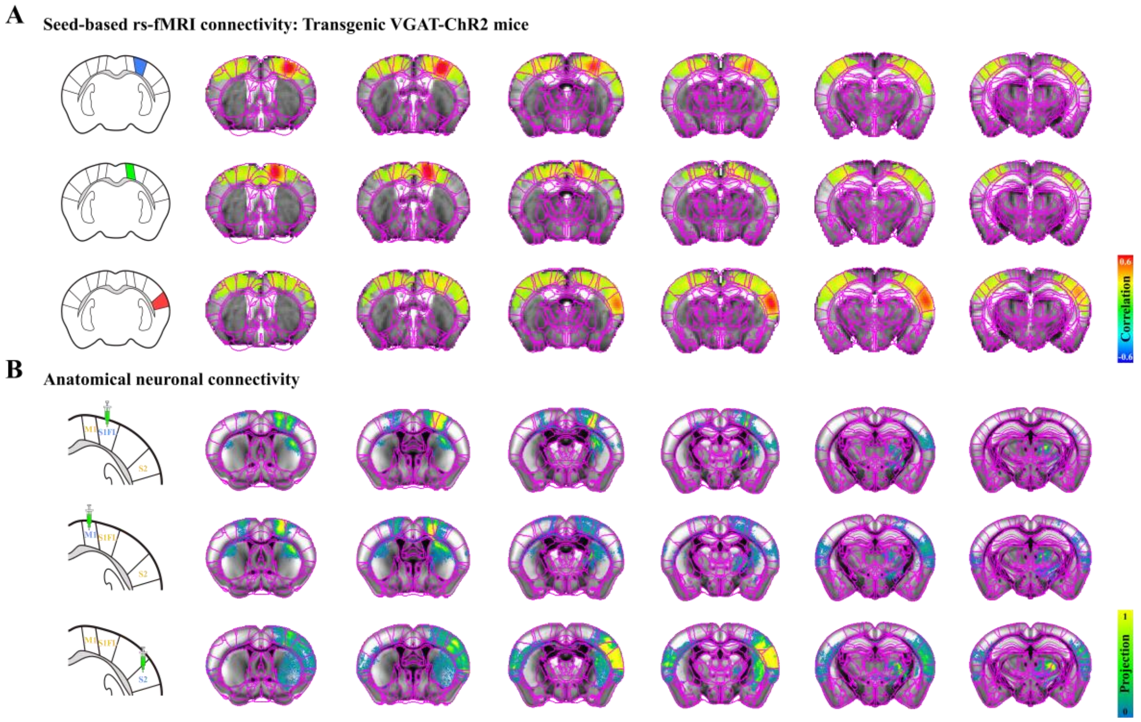


**Figure S4. Bilateral homotopic rs-fMRI connectivity is not primarily mediated by direct neural corticocortical connections.**

**(A)** Functional connectivity maps measured by seed-based rs-fMRI in VGAT-ChR2 mice. To determine whether bilateral homotopic correlation is a general feature of rs-fMRI independent of strain, functional connectivity was measured with the seed ROIs of S1FL, M1 and S2 in VGAT-ChR2 mice. Similar to the findings of naïve mice (Fig. 2B), resting-state fMRI in transgenic mice showed a strong correlation between bilateral somatosensory cortices but not between the cortex and thalamus in the ipsilateral hemisphere.

**(B)** Neural connectivity maps projected from somatosensory cortices, obtained from the Allen Institute (16). Ipsilateral projections in the somatosensory network are generally stronger than contralateral ones.

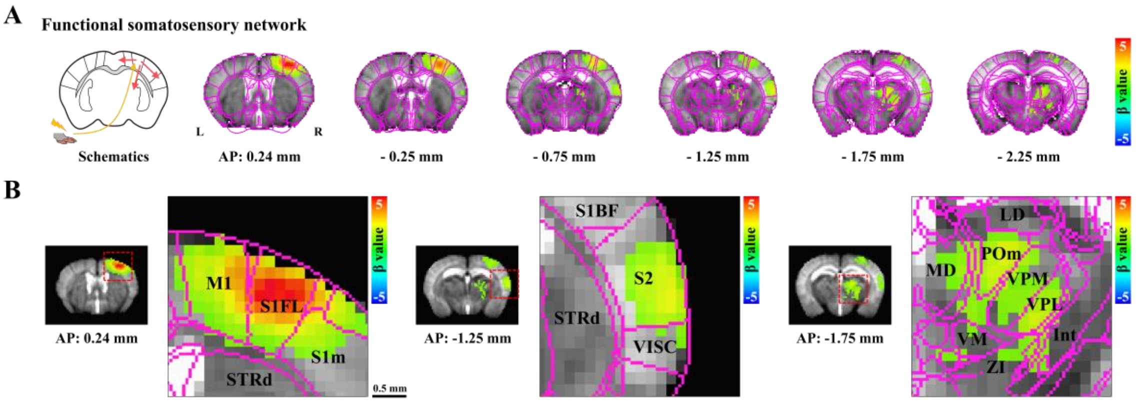


**Figure S5. Brain-wide fMRI maps of somatosensory-evoked activities.**

**(A)** Contralateral somatosensory regions, including the cortices (S1FL, M1 and S2) and thalamic nuclei (VPL and POm), responded to forepaw stimulation (VGAT-ChR2, n=22). Anterior-posterior slice coordinates relative to bregma are also marked. R, right; L, left hemisphere.

**(B)** Expanded fMRI maps in cortical and thalamic regions overlaid on the mouse brain atlas with labeling.

M1, primary motor area; S1FL, primary somatosensory area of forelimb; S1m, primary somatosensory area of mouth; STRd, striatum of dorsal region; S1BF, primary somatosensory area of barrel field; S2, secondary somatosensory area; VISC, visceral area; LD, lateral dorsal nucleus; MD, mediodorsal nucleus; POm, posterior medial nucleus; VPM, ventral posteromedial nucleus; VPL, ventral posterolateral nucleus; VM, ventral medial nucleus; Int, internal capsule; ZI, zona incerta; scale bar, 0.5 mm.

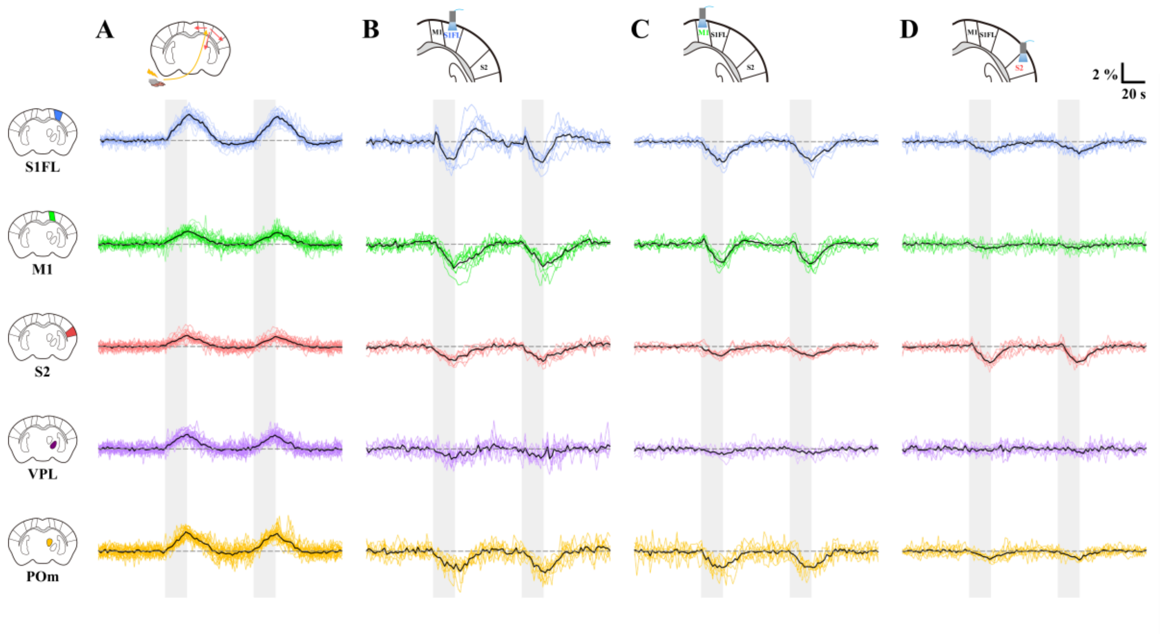


**Figure S6. CBV-weighted fMRI responses of individual animals in the somatosensory network during forepaw stimulation and optogenetic cortical inactivation.**

The fMRI time courses in the cortical (S1FL, blue lines; M1, green; S2, red) and thalamic (VPL, purple; POm, yellow) areas of individual animals during **(A)** forepaw stimulation, **(B)** S1FL inactivation, **(C)** M1 inactivation and **(D)** S2 inactivation were plotted. Black time course, animal-wise averaged response; gray vertical bar, 20-s stimulus.

One interesting observation is the CBV response at the optogenetically stimulating site: S1FL under S1FL Opto, M1 under M1 Opto, and S2 under S2 Opto. In the S1FL, an initial positive CBV response was followed by a negative CBV change and then a poststimulus positive CBV response. In S2, only negative CBV responses were observed. The initial positive CBV response was induced by activated inhibitory neurons, while the negative response was due to suppressed excitatory neurons. The underlying reason for different responses is unclear but could possibly be due to different combinations of active inhibitory vs. excitatory neurons in different regions. A large inter-animal variation was observed in the S1FL under S1FL Opto stimulation. This large variation was removed except for one animal when the difference between optogenetic stimulation with and without forepaw stimulation (Diff) was obtained. When we removed one outlier, the initial positive CBV response and the poststimulus overshoot disappeared in the averaged Diff time trace. However, these outlier data did not have much impact on an averaged trace (Fig. 4D $Diff_{S1FL}$).

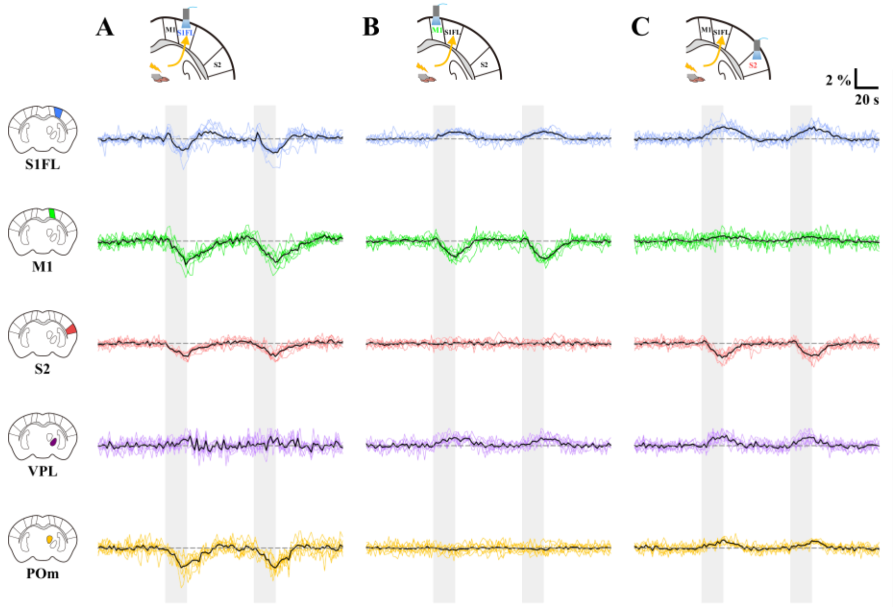


**Figure S7. CBV-weighted fMRI responses of individual animals in the somatosensory network during simultaneous cortical inactivation and forepaw stimulation.**

The fMRI time courses in the cortical (S1FL, blue lines; M1, green; S2, red) and thalamic (VPL, purple; POm, yellow) areas of individual animals during forepaw stimulation with **(A)** S1FL inactivation, **(B)** M1 inactivation and **(C)** S2 inactivation were plotted. Black time course, animal-wise averaged response; gray vertical bar, 20-s stimulus.

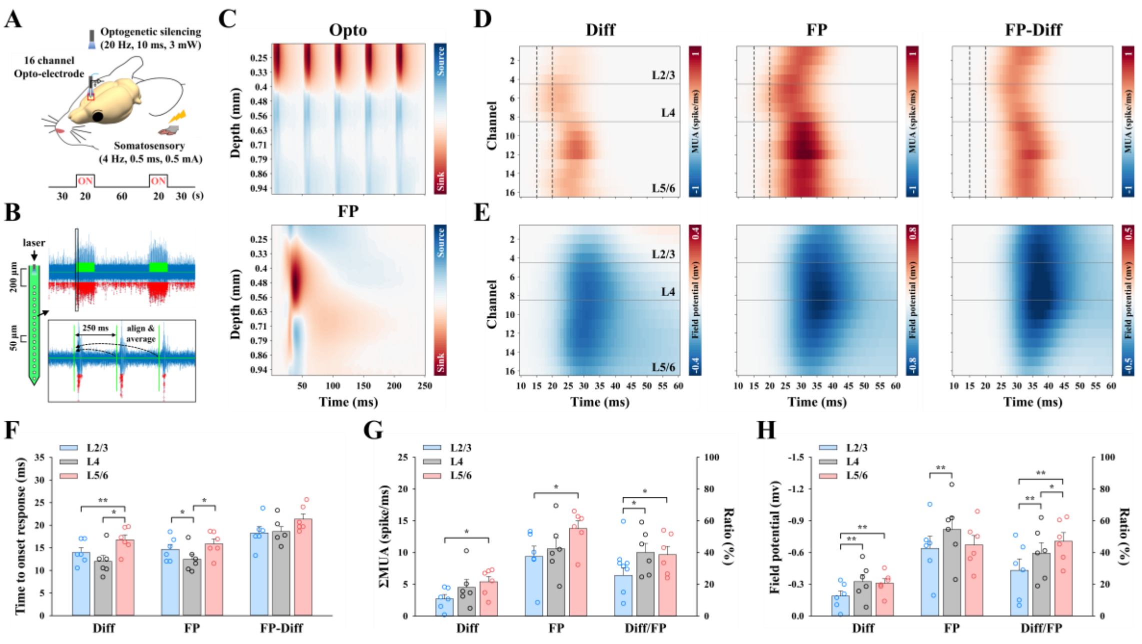

**Figure S8. Cortical depth-dependent MUA and LFPs to distinguish sensory input-driven activity and local recurrent activity.**

**(A)** Experimental schemes of electrophysiological recording with the 16-channel optoelectrode to measure somatosensory-evoked neural activity during S1FL inactivation (VGAT-ChR2, n=6). Optogenetic stimulation was performed on the surface of the S1FL cortex.

**(B)** Quantification of neuronal activity changes during stimulation. An example MUA trace with 4 Hz forepaw stimulation recorded from a single channel is shown (raw and expanded traces). The traces during two 20-s stimulus periods were aligned with a 250-ms window for comparison between neural activities acquired from 4 Hz somatosensory and 20 Hz optogenetic silencing. Spike activities from repeated MUA trains (or field potential from LFP waveforms) were then averaged across 20-s duration × 2 blocks × 4 windows/s × # of trials. Green line, forepaw stimulus duration; red dot, detected multiunit spike.

**(C)** Current source density (CSD) depth profile to identify the location of layer 4 (L4). Current sinks induced by optogenetic stimulation were found at depths of 0.2-0.4 mm below the cortical surface, whereas sink activity induced by forepaw stimulation was observed at depths of 0.4-0.56 mm (n=1), which is defined as L4. Red-to-blue, current sink-to-source.

**(D-E)** Time-dependent MUA and LFP depth profile in the S1FL (n=6) responding to somatosensory stimulation. **(D)** MUA trains and **(E)** LFP waveforms for the difference between optogenetic silencing and silencing/forepaw stimulation (Diff; input-driven) and for forepaw stimulation (FP) were compared to distinguish the local recurrent activity (FP-Diff). The earliest MUA response to forepaw stimulation occurred at L4, which is expected from thalamocortical inputs, and recurrent activity observed ~6 ms later. Gray lines, borders of the S1FL layer; black dashed lines in **(D)**, time periods of 15 and 20 ms after stimulus onset.

**(F)** Cortical depth-dependent neural onset time. Somatosensory input-driven MUA (Diff) was first evoked at L4 of the S1FL approximately 12 ms after stimulus onset, but local recurrent activity (FP-Diff) occurred similarly across the layers.

**(G-H)** Cortical depth-dependent neural strength and ratio of input-driven activity to total somatosensory-induced activity. **(G)** The MUA response peaked at L5/6, but **(H)** the LFP amplitude was stronger at L4. Thalamic input-driven activity was relatively lower in the upper cortical layers (blue bars) than in the granular and infragranular layers (gray and red bars). $^*p < 0.05$ and $^{**}p < 0.01$ (one-way analysis of variance with repeated measures followed by the Bonferroni *post hoc* test).

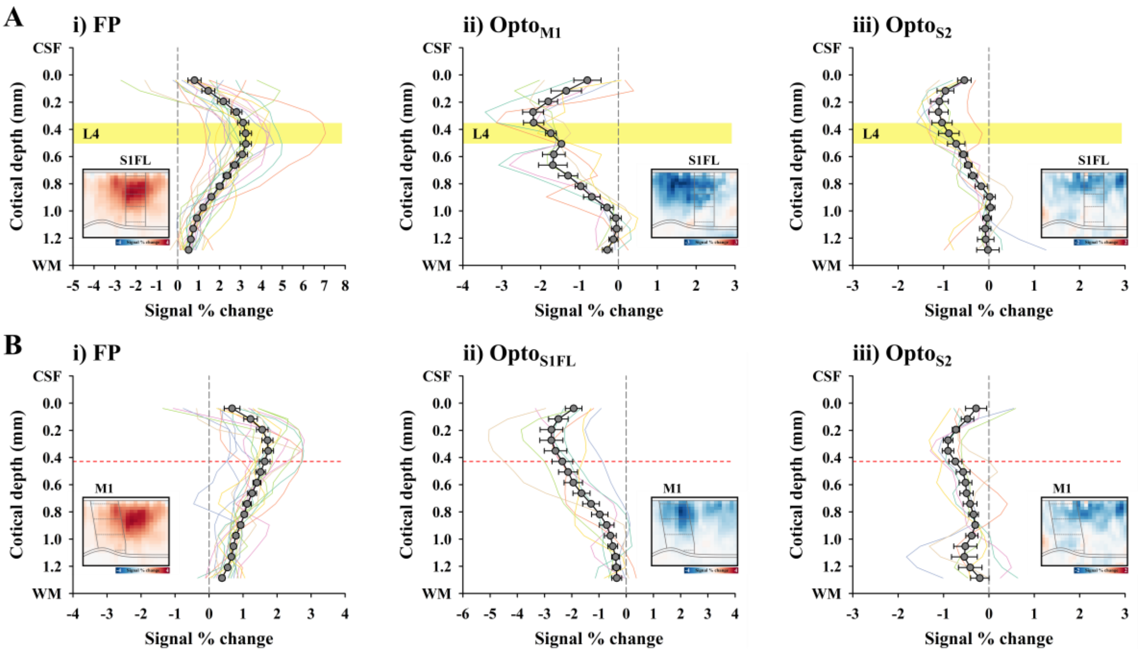


**Figure S9. Cortical depth-dependent CBV responses of individual animals in the S1FL and M1 during forepaw stimulation and optogenetic cortical inactivation.**

CBV responses in **(A)** the S1FL and **(B)** M1 of individual animals during forepaw stimulation and optogenetic cortical inactivation were plotted as a function of cortical depth. Inset images, averaged cortical-flattened fMRI maps; profiles with black circles, animal-wise averaged response; yellow-dashed box in A, ~150 µm width position of L4 of the S1FL; red-dashed line in B, laminar border of L2/3 and L5 in M1.

Although there was a large inter-animal variation, two peaks were observed in the S1FL when M1 was suppressed. When one animal profile with two strong peaks (green profile) was removed, the averaged profile of the remaining six animals also showed two peaks, indicating that the two peaks were indeed genuine and that the CBV point spread function was less than 150 µm FWHM.

**Table S1. Resting-state functional connectivity of the somatosensory network measured by resting-state fMRI and cortical inactivation fMRI.**

| ROI & Study | Downstream ROI | Ipsilateral hemisphere | | | | | Contralateral hemisphere | | | | |
|---|---|---|---|---|---|---|---|---|---|---|---|
| | | **S1FL** | **M1** | **S2** | **VPL** | **POm** | **S1FL** | **M1** | **S2** | **VPL** | **POm** |
| **S1FL** | **rs-fMRI (Z)** | **Seed-ROI** | 0.42±0.04 | 0.32±0.03 | -0.12±0.04 | -0.02±0.02 | 0.35±0.03 | 0.31±0.04 | 0.23±0.03 | -0.07±0.03 | -0.00±0.03 |
| | **Inactivation (%)** | **-1.41±0.19** | -1.34±0.17 | -1.03±0.07 | -0.65±0.05 | -1.29±0.11 | -0.31±0.03 | -0.21±0.05 | -0.05±0.05 | 0.09±0.08 | 0.10±0.11 |
| **M1** | **rs-fMRI (Z)** | 0.42±0.04 | **Seed-ROI** | 0.25±0.03 | -0.12±0.03 | 0.00±0.02 | 0.31±0.03 | 0.38±0.05 | 0.21±0.03 | -0.13±0.04 | -0.00±0.01 |
| | **Inactivation (%)** | -1.38±0.12 | **-1.29±0.15** | -0.71±0.06 | -0.33±0.08 | -1.29±0.17 | -0.28±0.07 | -0.45±0.10 | -0.08±0.09 | -0.04±0.15 | 0.13±0.05 |
| **S2** | **rs-fMRI (Z)** | 0.32±0.03 | 0.25±0.03 | **Seed-ROI** | -0.15±0.04 | -0.06±0.01 | 0.23±0.02 | 0.20±0.03 | 0.29±0.03 | -0.09±0.04 | 0.01±0.01 |
| | **Inactivation (%)** | -0.73±0.04 | -0.35±0.03 | **-1.04±0.08** | -0.13±0.05 | -0.50±0.07 | -0.11±0.05 | -0.11±0.05 | -0.31±0.03 | -0.06±0.06 | 0.03±0.22 |

The mean ± SEM is reported for resting-state fMRI (rs-fMRI, wild-type, 10 dataset from n = 5), S1FL inactivation (VGAT-ChR2, n = 8), M1 inactivation (VGAT-ChR2, n =7) and S2 inactivation (VGAT-ChR2, n = 7). The strengths of functional connectivity in rs-fMRI were calculated as the synchronization of resting fluctuations (z-score) with the seed-ROI area, whereas the spontaneous connectivity in cortical silencing fMRI was described as regional signal changes during optogenetic inactivation.

**Table S2. Multiunit activity characteristics of somatosensory-driven long-range input and local recurrent activity in the S1FL.**

| Measurement \ Study | Total somatosensory activity | Somatosensory input activity | Local recurrent activity |
|---|---|---|---|
| **Onset latency (ms)** | 14.83 ± 1.22 | 14.50 ± 1.67 | 20.00 ± 1.21**,## |
| **Peak latency (ms)** | 30.83 ± 1.40 | 26.50 ± 1.43** | 32.83 ± 1.45**,## |
| **FWHM** | 14.41 ± 0.88 | 12.36 ± 0.94** | 12.55 ± 0.96* |
| **Peak amplitude (spike/ms)** | 0.79 ± 0.11 | 0.37 ± 0.07** | 0.59 ± 0.90* |
| **∑MUA (spike/ms)** | 11.71 ± 1.34 | 4.55 ± 0.81** | 7.45 ± 0.92** |
| **Goodness of fit ($R^2$)** | > 0.99 | > 0.99 | > 0.99 |

The mean ± SEM is reported for the 16-channel averaged multiunit activity (MUA) of the total somatosensory activity, input activity, and local recurrent activity (VGAT-ChR2, n = 6). The dynamic properties, including latencies (times to onset and peak of response) and full width at half-maximum (FWHM), and the MUA response, including peak amplitude and ∑MUA, were determined from the MUA trace fitted with a single Gaussian function. Onset latency was defined as the time for the first bin of five continuous bins whose spike rate differed significantly from the prestimulus baseline spike rate ($p < 0.05$, one-sample t-test). The goodness-of-fit between the single Gaussian function and the MUA trace is presented as the $R^2$ value. *$p < 0.05$ and **$p < 0.01$ for vs. total somatosensory activity, $^{\#}p < 0.05$ and $^{\#\#}p < 0.01$ for vs. somatosensory input activity (one-way analysis of variance with repeated measures followed by the Bonferroni *post hoc* test).

**Table S3. Local field potential characteristics of somatosensory-driven long-range input and local recurrent activity in the S1FL.**

| Measurement \ Study | Total somatosensory activity | Somatosensory input activity | Local recurrent activity |
|---|---|---|---|
| **Peak latency (ms)** | 36.00 ± 1.77 | 31.67 ± 1.52** | 38.00 ± 1.71**,## |
| **FWHM** | 20.62 ± 1.21 | 19.38 ± 1.12** | 18.59 ± 1.62* |
| **Peak amplitude (mv)** | -0.69 ± 0.10 | -0.28 ± 0.05* | -0.46 ± 0.09* |
| **∑LFP (mv)** | -23.10 ± 3.30 | -7.36 ± 1.31* | -16.30 ± 3.34* |
| **Goodness of fit ($R^2$)** | > 0.99 | 0.99 ± 0.01 | 0.99 ± 0.00 |

The mean ± SEM is reported for the 16-channel averaged local field potential (LFP) of the total somatosensory activity, input activity, and local recurrent activity (VGAT-ChR2, n = 6). The dynamic properties, including peak latency and full width at half-maximum (FWHM), and the MUA response, including peak amplitude and ∑LFP, were determined from the LFP waveform fitted with a double gamma variate function. The goodness-of-fit between the gamma variate function and the LFP waveform is presented as the $R^2$ value. *$p < 0.05$ and **$p < 0.01$ for vs. total somatosensory activity, #$p < 0.05$ and ##$p < 0.01$ for vs. somatosensory input activity (one-way analysis of variance with repeated measures followed by the Bonferroni *post hoc* test).
